# Pomegranate: 2D segmentation and 3D reconstruction for fission yeast and other radially symmetric cells

**DOI:** 10.1101/2020.07.07.191932

**Authors:** Erod Keaton Baybay, Eric Esposito, Silke Hauf

## Abstract

Three-dimensional (3D) segmentation of cells in microscopy images is crucial to accurately capture signals that extend across optical sections. Using brightfield images for segmentation has the advantage of being minimally phototoxic and leaving all other channels available for signals of interest. However, brightfield images only readily provide information for two-dimensional (2D) segmentation. In radially symmetric cells, such as fission yeast and many bacteria, this 2D segmentation can be computationally extruded into the third dimension. However, current methods typically make the simplifying assumption that cells are straight rods. Here, we report Pomegranate, a pipeline that performs the extrusion into 3D using spheres placed along the topological skeletons of the 2D-segmented regions. The diameter of these spheres adapts to the cell diameter at each position. Thus, Pomegranate accurately represents radially symmetric cells in 3D even if cell diameter varies and regardless of whether a cell is straight, bent or curved. We have tested Pomegranate on fission yeast and demonstrate its ability to 3D segment wild-type cells as well as classical size and shape mutants. The pipeline is available as macro for the open-source image analysis software Fiji/ImageJ. 2D segmentations created within or outside Pomegranate can serve as input, thus making this a valuable extension to the image analysis portfolio already available for fission yeast and other radially symmetric cell types.

## Introduction

Microscopy is a powerful technique to collect spatial and temporal information on cellular processes. However, collecting quantitative data from microscopy images represents a persistent bottleneck. This task is greatly aided by automated or semi-automated image analysis pipelines^1-3^. Automation minimizes subjectivity in the analysis, saves time, and allows for the quantification of more cells than manual analysis. This increases statistical power and provides fine-grained information on cell-to-cell variability.

Regions of interest (ROIs) can be delineated either in two dimensions (X/Y, 2D) or three dimensions (X/Y/Z, 3D). Analysis in 3D becomes important when objects of interest stretch across more than one optical section. Different strategies have been used to construct 3D ROIs. A common approach is the use of a fluorescent marker to segment objects of interest (e.g. nuclei or cells). Segmentation can be achieved by traditional image processing algorithms such as adaptive thresholding and binary morphometric operations, or by machine learning or deep learning techniques^4-14^. However, any strategy that uses a fluorescent marker for live-cell segmentation (i) increases phototoxicity, (ii) limits the number of fluorescent channels available for proteins or structures of interest, and (iii) can cause variability in segmentation when uneven illumination, or uneven staining or expression, lead to variability in fluorescence signal.

Using brightfield images to segment cells largely eliminates these problems but comes with its own set of challenges. While brightfield microscopy highlights the cell contour which allows for 2D segmentation, 3D information on cell shape is not readily available^15,16^. Recent efforts have begun to train neural networks to extract 3D information from brightfield Z-stacks by co-training with fluorescence markers^17^, but feasibility for whole-cell segmentation has not yet been demonstrated. Hence, additional methods are currently required to extend a 2D cell ROI obtained by brightfield imaging into 3D. When the cells being imaged are radially symmetric (e.g. rod-shaped), this additional information can be used to infer the cellular shape in 3D from that in 2D. Examples of radially symmetric cells include many types of bacteria (e.g. *E. coli* or *B. subtilis*^18-20^) and fungi (e.g. fission yeast^21-24^).

Fission yeast *(Schizosaccharomyces pombe*) is a eukaryotic model organism frequently used in single-cell imaging studies to gain an understanding of the cell cycle, cell polarity, cell division, and other intracellular signaling events^25,26^. Features that contribute to fission yeast’s popularity include its well-annotated, compact genome^27,28^, excellent molecular genetics tools^29^, and the presence of many genes and pathways that are shared with animals and plants^30,31^. A number of segmentation pipelines based on brightfield imaging have been developed specifically for fission yeast or other radially symmetric cells but only perform 2D segmentation^32-36^.

Here, we present Pomegranate, a pipeline for the 3D segmentation of radially symmetric cells which combines 2D brightfield segmentation with 3D morphological extrapolation. We validate Pomegranate using fission yeast. Unlike previous extrapolation approaches (e.g.^37,38^), Pomegranate does not assume cells to be straight, nor does it assume constant cell width. To create the 3D shape, Pomegranate uses spheres with centers located along the central axis of the cell. The diameter of each sphere is taken from the cell width at this position. Thus, Pomegranate accurately projects the variable cell width of a single cell into 3D and performs equally well on bent, curved and straight cells. Pomegranate is expected to work on all cell types that are radially symmetric along a central axis that is parallel to the imaging plane and provides improved 3D segmentation for such cells. Pomegranate is available as a macro for the open-source image analysis software FIJI/ImageJ.

## Results

### 3D reconstruction with Pomegranate

We developed Pomegranate to segment fission yeast or other radially symmetric cells in 3D. As input, the pipeline takes a Z-stack of brightfield images for segmenting the cell and a Z-stack of fluorescence images for segmenting the cell nucleus and any markers of interest (Fig. 1). Before being added to the pipeline, the images are background-subtracted, flat-field corrected and corrected for axial chromatic aberration as well as for distortions in axial distance (Fig. S1 and S2).

**Figure 1.**
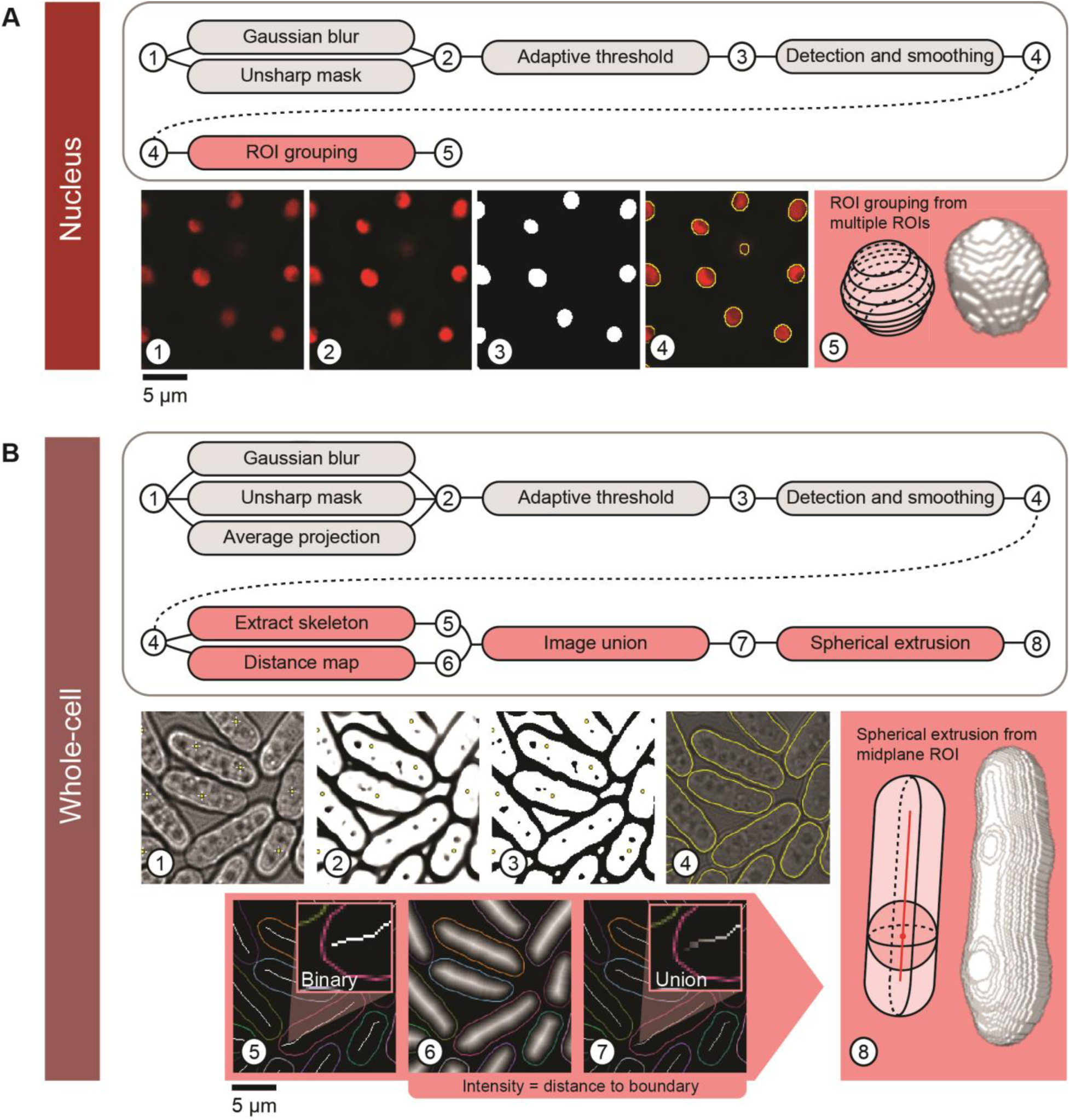
Pomegranate pipeline schematic. **(A)** Nuclear segmentation and reconstruction - (Top) Schematic of processing steps for 2D nuclear segmentation (grey) and 3D reconstruction (red). (Bottom) Sequential images through the segmentation and reconstruction process: (1) Sample slice of an input image, showing nuclear marker TetR-tdTomato-NLS, (2) sample slice of a processed input image following Gaussian blur and unsharp mask, (3) sample slice of a binary image from adaptive thresholding, (4) smoothed ROIs derived from binary image, (5) cartoon of reconstruction method and representative model from 3D reconstruction as a result of ROI grouping. **(B)** Whole-cell segmentation and reconstruction - (Top) Schematic of processing steps for 2D whole-cell segmentation (grey) and 3D reconstruction (red). (Bottom) Sequential images through the segmentation and reconstruction process: (1) Sample slice of an input brightfield image with point selection on nuclear centroids from A (for the optional Z-alignment of whole-cell reconstructions to the corresponding nuclei). (2) Z-projection of multiple slices processed with Gaussian blur and unsharp mask, (3) binarization of projection, (4) smoothed ROIs derived from binary image (yellow) overlaid on brightfield image, (5) topological skeleton of binary image, (6) Euclidean distance map of binary, (7) union of topological skeleton and Euclidean distance map, used as radius profile input for spherical extrusion when generating the whole-cell 3D reconstruction, (8) cartoon of reconstruction method and representative model from 3D reconstruction as a result of spherical extrusion. For simplicity, extrusion is shown here along a straight axis, but topological skeletons are typically not straight (panels 5,7).

Nuclear segmentation is optional in Pomegranate, and nuclei and cells are initially treated separately and later combined. To define the nucleus, Pomegranate uses traditional adaptive thresholding across all optical sections (Fig. 1A, see Methods for details). The ROIs from all optical sections are combined to obtain the 3D representation of the nucleus. We used TetR-tdTomato-NLS as the fluorescent marker, which yields a bright signal in the nucleus and a slightly weaker signal in the nucleolus^39^.

For whole-cell segmentation, Pomegranate uses brightfield images to define a midplane ROI (Fig. 1B, see Methods for details). To obtain the midplane ROI, the characteristic intensity pattern surrounding cells in the brightfield image (Fig. S3) is used to define the edges of the cell^33,35,40^ (Fig. 1B, see Methods for details). As an alternative to this brightfield segmentation, externally prepared midplane ROIs can be used. This allows users to run any pipeline of their choice for 2D cell segmentation (e.g.^33,34,41^) while still profiting from Pomegranate’s 3D reconstruction. We demonstrate this by using 2D brightfield segmentation obtained by “YeaZ”^41^, a convolutional neural network (CNN) trained on yeast brightfield images (Fig. S4A). YeaZ-segmented ROIs were imported into Pomegranate and successfully used in place of Pomegranate midplane ROIs to produce 3D representations of cells (Fig. S4A). While the accuracy of 2D segmentation between Pomegranate and YeaZ was comparable (Fig. S4B), the accuracy of YeaZ segmentation could potentially exceed that of Pomegranate with additional model training.

The key feature of Pomegranate is extrusion of the cellular midplane ROI into 3D (Fig. 2). Based on the assumption of radial symmetry, Pomegranate extrudes multiple spheres along the length of the cell, whose diameters correspond to the cell width at each position (Fig. 2A-D). ROIs in the optical sections away from the midplane are defined by this extrusion (Fig. 2B,D). The ROIs from all optical sections are then combined to obtain the 3D representation of the cell (Fig. 2E,F). As path for the centers of the spheres, Pomegranate uses the ‘topological skeleton’ (Fig. 1B, panel 5 and 7; Fig. 2C), a line equidistant to the cell boundaries. Since this line follows cell shape, Pomegranate processes straight, bent or curved cells equally well.

**Figure 2.**
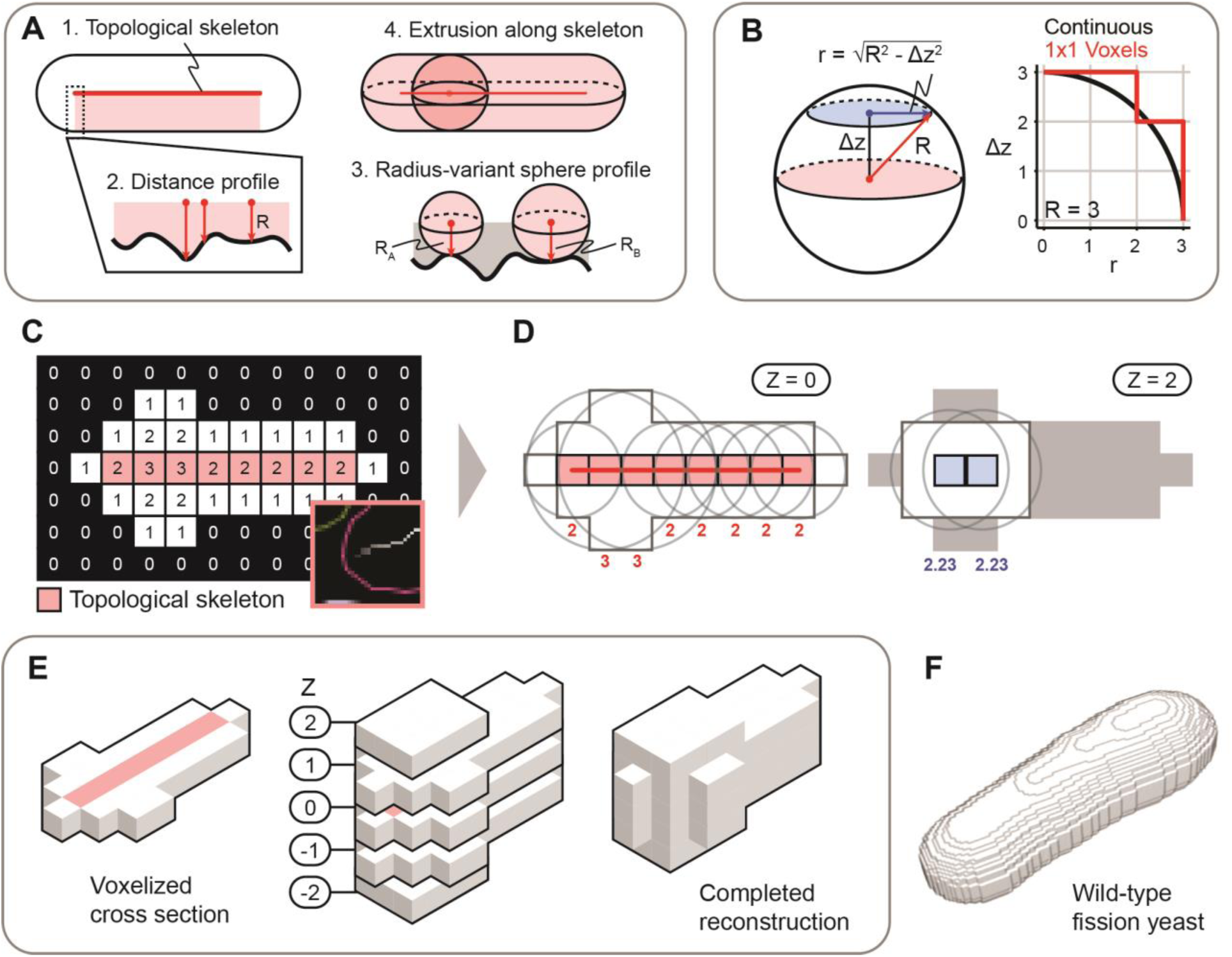
Whole-cell reconstruction algorithm. **(A)** General schematic for the whole-cell 3D reconstruction algorithm: From a 2D binary image in the midplane of a cell, spheres are extruded along the topological skeleton, where the radii of these spheres is the distance to the nearest edge of the cell. Note that the topological skeleton is shown here as a straight line but does not have to be straight. **(B)** At each optical section away from the midplane (distance: Δ z), a voxel is included in the reconstruction when the majority of its volume is part of the sphere profile. (Right) Black line represents the sphere profile, red line represents the voxelized z-section as seen in C. **(C)** Coarse 2D image converted into a distance map where the values in each pixel represent the Euclidean distance to the periphery of the binary, with the topological skeleton annotated in red. Inset: example of union between topological skeleton and distance map, from Figure 1B, panel 7. The greyscale of the pixels of the topological skeleton indicates distance to the periphery. **(D)** Sample points on topological skeleton at Z = 0 (left) and Z = 2 (right) with example spherical cross sections centered in XY along the topological skeleton. Cross section radii are determined by the equation described in (B). Radii are indicated in red at Z = 0 and blue at Z = 2. **(E)** Visualization of the coarse reconstruction in 3D, with all optical sections of the complete reconstruction shown. **(F)** Example 3D reconstruction of a wild-type cell (voxel size 0.1071 μm x 0.1071 μm x 0.065 μm).

Finally, if both cell and nuclear segmentation are available, nuclei are assigned to the cells in which they are located. If the Z positions of the cellular and nuclear midplane differ, users have the option to re-assign cellular midplane ROIs to the optical section of the corresponding nuclear midplane. This may be necessary if cells are not all located in the exact same plane.

Overall, the core unique step of Pomegranate is the spherical extrusion to obtain the 3D representation of cells (Fig. 2). Beyond that, the pipeline is adaptable to the specific requirements of each individual user.

### Pomegranate accurately isolates regions of interest

To assess the ability of Pomegranate to accurately segment cells, we applied the pipeline to cells expressing the nuclear fluorescent marker TetR-tdTomato-NLS in combination with a cytoplasmic fluorescent marker: Mep33-mCherry for a weaker cytoplasmic signal or Mep33-tdTomato for a brighter cytoplasmic signal (Fig. 3A). We used the weaker cytoplasmic signal when testing 3D nuclear segmentation and the brighter cytoplasmic signal when testing the ability of the 3D cellular segmentation to capture cytoplasmic signals. After segmentation and 3D reconstruction, we analyzed the intensities of voxels either in the full image, or within the ROIs defining nuclei or cells. Pomegranate successfully isolated the higher intensity signals in nuclei or cells from the background signal (Fig. 3B), indicating that Pomegranate segments accurately in 3D.

**Figure 3.**
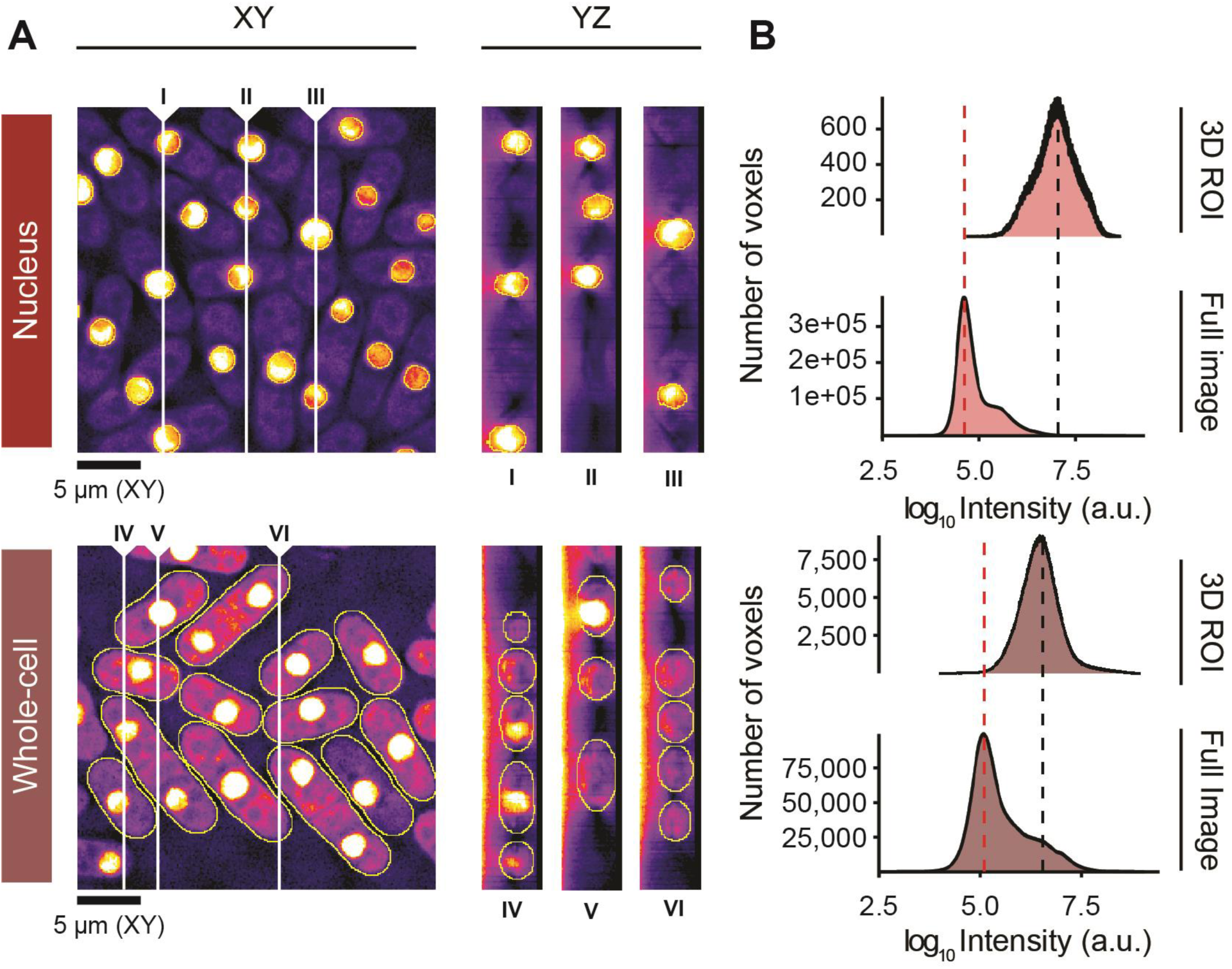
Signal acquisition performance. **(A)** Representative optical sections in XY and three sample YZ orthogonal views at different X positions. White vertical lines indicate sampling positions for corresponding YZ orthogonal views. Voxel size is 0.1071 μm x 0.1071 μm x 0.065 μm. Images are from fission yeast expressing TetR-tdTomato-NLS and Mep33-mCherry for nuclear analysis (top) and TetR-tdTomato-NLS and Mep33-tdTomato for whole-cell analysis (bottom). Annotations provide minimum and maximum intensities. All images have been subject to gamma correction (γ = 0.50) to improve visibility of cytoplasmic signal. **(B)** Intensity histograms for all voxels in the image (Full Image), and for all voxels contained within the 3D ROIs. Dashed lines annotate maxima: background (red) and signal of interest (black).

A frequent workaround to proper 3D segmentation is applying the 2D midplane ROI to all optical sections, which is analogous to analyzing a projection of all optical sections^42-44^ (Fig. S5A). This strategy falsely includes extracellular background. The approach can be slightly improved by only including sections above or below the midplane that are likely to be located within the cell^45^ (Fig. S5A). Direct comparison of this approach to Pomegranate finds that Pomegranate identifies a narrower distribution of pixel intensities (Fig. S5B), likely reflecting a more accurate capture of cytoplasmic signals. We conclude that the brightfield-based 3D segmentation of Pomegranate is successful in capturing fluorescent cytoplasmic signals and does so with higher accuracy than projection-based approaches.

### Pomegranate refines volume measurement in wild-type cells

Segmenting cells in 3D is useful to measure cellular signals but can also provide information on the volume and surface area of cells. Both of these size parameters are important in cell growth studies^46,47^. To estimate cell size, some prior studies have made the approximation that fission yeast cells are cylinders with hemispherical ends (e.g.^37,38,48,49^). Cell length and width measurements from the midplane ROI are used to calculate the volume of this idealized shape. To compare this method to 3D reconstruction by Pomegranate, we imaged wild-type fission yeast cells and analyzed more than 8,000 cells with two variations of this earlier method in addition to Pomegranate. In the two variations, cell length and width are obtained either from the major and minor axis of an elliptical fit or from the maximum and minimum Feret diameter (also called the caliper diameter) of the midplane ROI (Fig. 4A, S6, see Methods for details). We find that cellular volumes obtained by Pomegranate are around 20 % smaller (Fig. 4B). Volumes obtained by Pomegranate are also less variable across the cell population (Fig. 4B). This suggests that the methods assuming a straight rod shape tend to overestimate volume and that the Pomegranate ROI more accurately represents cell shape. The overlay of example regions shows that the discrepancies largely result from unequal width along the length of the cells (Fig. 4C). In addition, the requirement of the elliptical fit to maintain the area of the original ROI tends to overestimate cell length (Fig. 4C). Surface area was slightly larger when measured by Pomegranate when compared to the surface areas calculated from the idealized rod-shape (Fig. S6B), possibly because Pomegranate better captures some of the irregularity in fission yeast’s whole-cell morphology. Altogether, this analysis suggests that 3D segmentation by Pomegranate improves volume and surface measurements compared to methods that assume a cylinder with hemispherical ends as cell shape.

**Figure 4.**
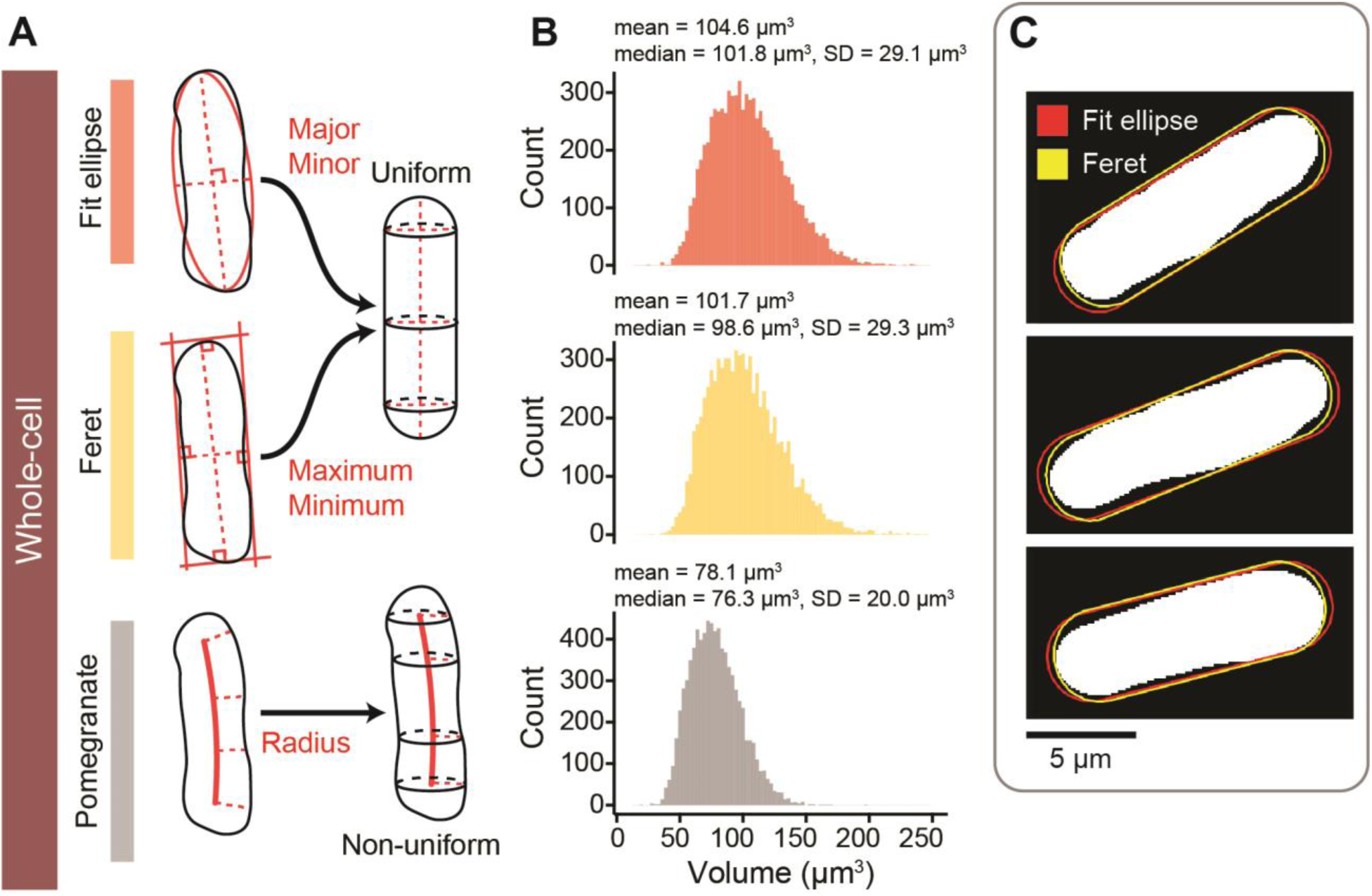
Comparison of methods for calculating wild-type cell volume. **(A)** Schematic representations of methods for calculating whole-cell volume. Parameters are extracted from the cell midplane ROI (left) and used in generating 3D volume (right). The ‘Fit ellipse’ method extracts the major and minor axes of an ellipsoidal fit as length and width, the ‘Feret’ method extracts the maximum and minimum Feret diameter as length and width. These are used as input to generate a rod-shaped cell. Pomegranate generates a non-uniform volume as described in Figure 2. **(B)** Histograms (binwidth = 3; Freedman-Diaconis) of cell volumes from the same population of 8,026 cells, calculated with the three approaches: Fit Ellipse method (red); Feret method (yellow); and Pomegranate (grey). Annotations provide descriptive statistics (SD = standard deviation). **(C)** Midplane ROI (white), with overlaid cross sections of the rod-shape obtained with length and width from ellipse fitting (red) or Feret diameters (yellow). Three representative cells are shown.

The volume of the nucleus scales with the volume of the cell^37,50^ (Fig. S7). In fission yeast, the ratio of nuclear to cell volume (the N/C ratio) has been reported to be 8 % when using idealized geometries for both nucleus and cell^37,49^. Electron microscopy experiments measured 9.4 %^51^ and 12 %^52^. Similarly, we find an N/C ratio of 10 % (SD = 2.4 %) when using volumes from idealized geometries (from Feret diameters, Fig. S7B). Measured by Pomegranate 3D reconstruction, the N/C ratio is higher with a value of 12.7 % (SD = 2.8 %, Fig. S7B), as is expected from the smaller cell volume (Fig. 4B). We also find that the N/C ratio is more variable in smaller compared to larger cells (Fig. S7C). This suggests that cell division introduces variability, which may get corrected as cells grow^49,53^.

Given the difficulties in measuring volume, cell length has sometimes been used as a simple proxy for cell size (e.g.^54,55^). Correlation analysis in our large dataset (Fig. S8) shows that cell volume indeed correlates well with cell length (Pomegranate cell volume and Feret length, Pearson r = 0.90, p < 0.01). However, cell volume also correlates to a lesser yet significant extent with cell width (Pomegranate cell volume and Feret width, Pearson r = 0.60, p < 0.01). The correlation between cell length and cell width is weaker (Feret length and Feret width, Pearson r = 0.28, p < 0.01). This suggests that cell width, which is often ignored, influences cell volume largely independent of cell length and should be accounted for in order to approximate volume. The analysis above further suggests that Pomegranate’s ability to take the variation of cell width within individual cells into account improves cell volume estimates.

### Pomegranate reconstructs cell size and shape mutants in 3D

While wild-type fission yeast cells have a relatively uniform shape, mutant cells can drastically deviate from this shape^56^. We therefore tested whether Pomegranate can segment cell size and shape mutants. We used *tea1Δ* for curved cells, *wee1-50* for short cells, and *cdc25-22* for long cells^57-59^. Our whole-cell segmentation successfully captured the shape of these mutants (Fig. 5B). We estimated curvature by assessing solidity of the midplane ROI (Fig. 5A). Solidity is defined as the ROI area divided by the smallest convex area that contains the ROI (convex hull). Low solidity indicates a concave shape. Wild type and *tea1Δ* cells exhibited mean solidities of 0.94 and 0.93, respectively. The coefficient of variation for *tea1Δ* (CV = SD/Mean = 3 %) was higher than that of wild type (CV = 2 %), suggesting greater variability in cell curvature^57,60^. The *cdc25-22* cells exhibited the lowest mean solidity of 0.87. Because these cells are overly long, even slight curvature leads to large area differences between convex hull and ROI area. The highest solidity was exhibited by *wee1-50* mutant cells (0.95) as their compact geometry prohibits curvature. Our results are highly similar to those obtained by Liu *et al*. in the testing their morphometry toolbox^34^. This validates our 2D segmentation and demonstrates that Pomegranate successfully segments size and shape mutants.

**Figure 5.**
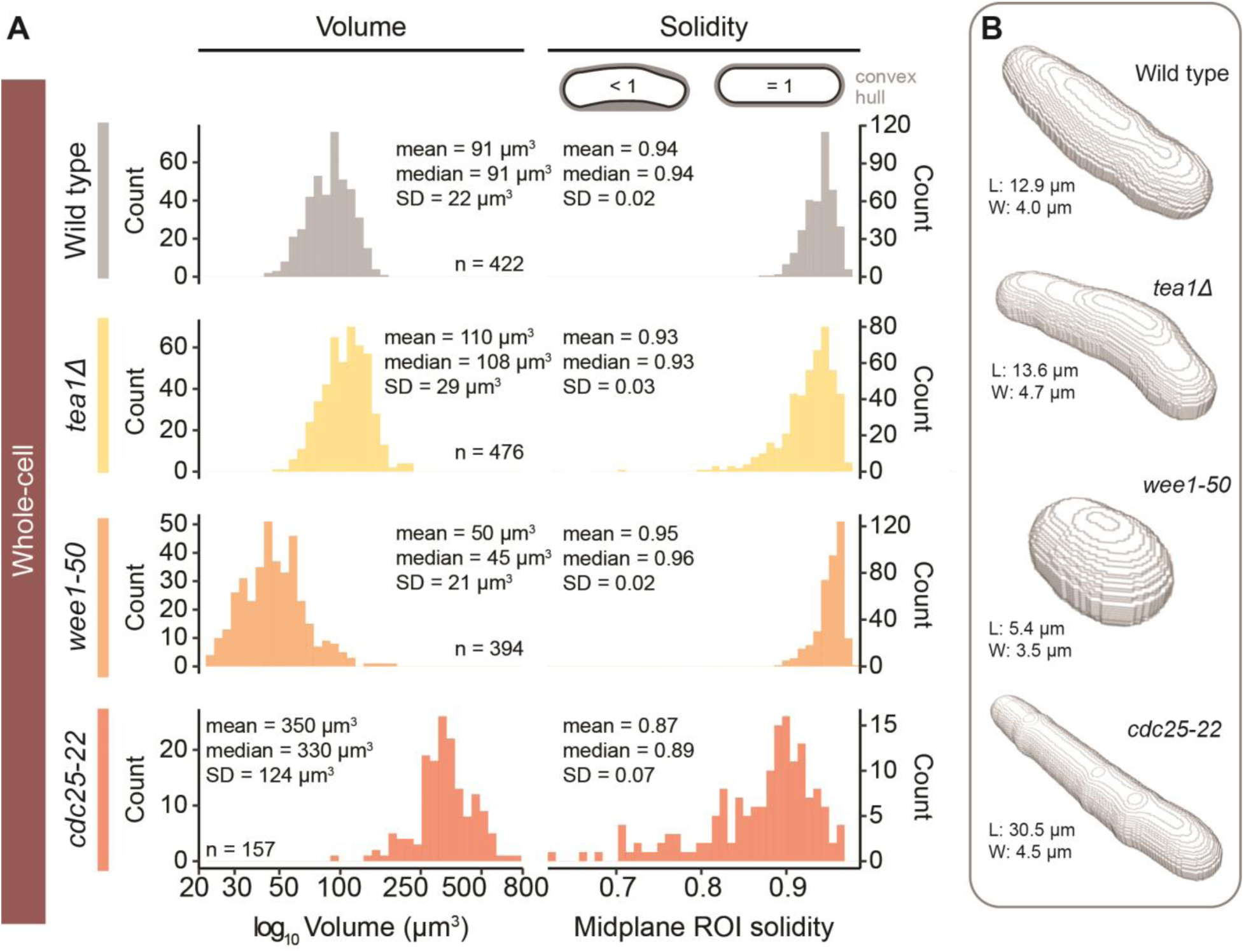
Size and shape of mutant strains captured by Pomegranate 3D reconstruction. **(A)** Histograms (bins = 40) of volume (left) and midplane ROI solidity (right), representing size and curvature, respectively, for wild type and cell shape and cell size mutants. The x-axis of the volume histogram is on a base-10 logarithmic scale. Annotations provide descriptive statistics (SD = standard deviation). **(B)** Reconstruction models of representative, individual cells. Annotations provide size parameters for each cell to illustrate scale: maximum Feret diameter (length, L) and minimum Feret diameter (width, W).

As the curvature analysis indicates, these size and shape mutants are not well approximated by assuming cells to be a cylinder with hemispherical ends (Fig. 4A). Pomegranate therefore provides the necessary functionality to measure volume of these mutants (Fig 5). Compared to wild-type cells with a mean volume of 91 μm^3^, *wee1-50* mutants, as expected, showed a smaller mean volume (50 μm^3^), and *cdc25-22* mutants a larger one (350 μm^3^). The mean volume of *tea1Δ* cells (110 μm^3^) was slightly larger than for wild-type cells, consistent with a slightly longer length at division than wild-type cells^61^. When we assumed an idealized rod shape instead, the volumes of the size and shape mutants were up to about 75 % larger than those obtained from the Pomegranate analysis (e.g. *cdc25-*22 Mean = 622 μm^3^, Fig. S9). This suggests that the advantages of using Pomegranate for 3D segmentation and volume estimation are even greater when analyzing size and shape mutants.

In summary, we conclude that Pomegranate successfully segments wild-type fission yeast cells as well as size and shape mutants in three dimensions (Fig. 3-5) and allows for accurate estimates of volume (Fig. 4). After correcting for axial chromatic aberration, the ROIs across all optical sections can be used to quantify any signal of interest (Fig. 6). Pomegranate is expected to work for all radially symmetric cells that can be sufficiently resolved and whose symmetry axis is parallel to the imaging plane. We show examples of bacteria in Fig. S10.

**Figure 6.**
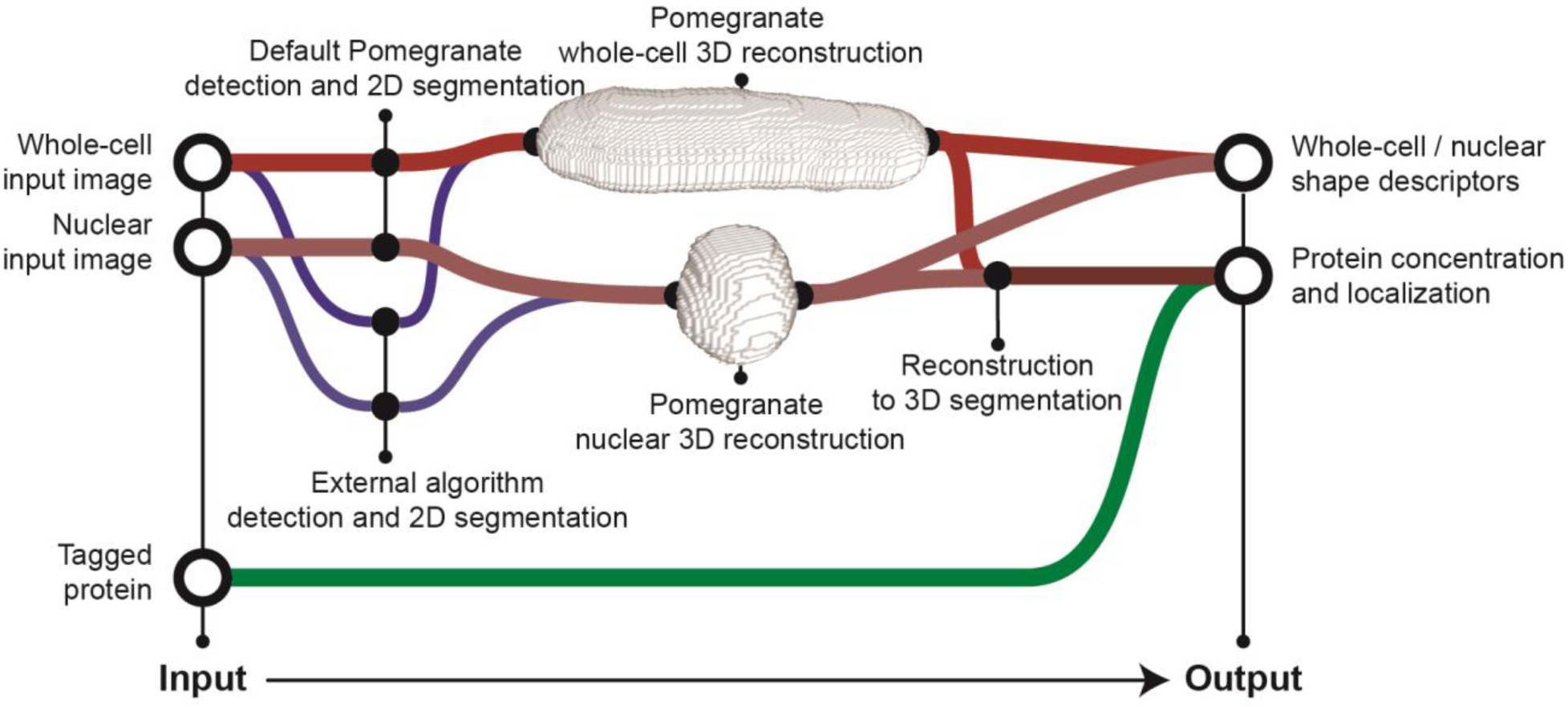
Workflow schematic. General schematic of pipeline input and output. Reconstructions are not to scale. For detection and 2D segmentation, either Pomegranate (red lines) or an external algorithm (blue lines) can be used. For Pomegranate 2D segmentation, a brightfield Z-stack is required as whole-cell input image, a fluorescence Z-stack is required as nuclear input image. After reconstruction and 3D segmentation, any desired signal that was co-recorded (“Tagged protein”) can be quantified.

## Discussion

### Different approaches to 2D segmentation from brightfield images

Using brightfield images for segmentation is appealing because acquiring these images is minimally phototoxic and all fluorescence channels remain available for proteins of interest. Brightfield microscopy yields a consistent intensity pattern along the cell boundary across optical sections (Fig. S3), which we and others have leveraged to obtain a midplane ROI. For example, PombeX^35^ identifies this pattern by k-means clustering. MAARs^33^ and the MPCS algorithm^32^ use correlation analysis. These methods have been further complemented by machine learning^35,62^ or neural networks^36^. Other methods do not use the pattern itself, but use traditional segmentation on brightfield images: either edge detection^63^ or adaptive thresholding^34^. Our segmentation relies on projection of the pattern and adaptive thresholding supplemented by watershedding and traditional morphometry tactics, which together have the advantage of simplicity, low computational demand, and no need for training data.

### Optional user input to Pomegranate

To accommodate different imaging conditions, several parameters in the 2D segmentation of Pomegranate (Fig. 1) can be changed by the user as specified in the usage guide available on GitHub. For the settings we used, up to 96 % of cells were accurately detected on images with low cell density. This percentage tended to be lower on high cell density images (between 70 and > 90 %). Adjustable filters exclude most false-positive ROIs (such as those capturing background or debris rather than a cell). Any remaining false-positive ROIs can be manually excluded prior to the 3D reconstruction step.

Sensitivity of image recognition algorithms to the imaging conditions is a general problem, and we have therefore constructed Pomegranate to be capable of using ROIs obtained by other segmentation methods as input for 3D reconstruction. These segmentations do not need to be performed on a brightfield image. Hence, 3D reconstructions with Pomegranate can also be performed from phase contrast imaging (Fig. S10), differential interference contrast imaging, widefield fluorescence microscopy, or confocal microscopy, provided a 2D detection and segmentation algorithm produces an accurate 2D cell shape as input.

### Advantages of 3D segmentation by Pomegranate

Because segmentation in 3D is not trivial, workarounds have often been used. Prominent workarounds are projections^42-44^ (Fig. S5) or approximations of the rod shape as a cylinder with hemispherical ends^37,38,48,49^ (Fig. 4). Both of these approaches require a 2D midplane ROI. Pomegranate uses the same midplane ROI but makes optimal use of the information contained in this ROI for 3D extrusion, thereby achieving a more accurate representation of the cell compared to these approaches.

Our analysis indicated that fission yeast cell width varies along the length of a cell (Fig. 4) and among cells (Fig. S8) and should not be ignored when calculating volume or representing cells in 3D. For the same reason, recent extrusion approaches to measure cell volume and surface area have already started to use techniques that take variations of cell width along the length of the cell into account^64-66^. Pomegranate differs from these methods in that it uses the topological skeleton rather than a straight central axis for extrusion. Pomegranate therefore more accurately represents bent or curved cells in 3D, and particularly excels on shape mutants (Fig. S9). Pomegranate provides 3D segmentation within the open-source image analysis software Fiji^3^ so that signals of interest in other channels can be easily quantified.

### Pipeline limitations

Pomegranate’s approach to 3D reconstruction still makes several simplifying assumptions. The spherical extrusion assumes that cells are indeed radially symmetric around the long axis, which may not always be the case. For fission yeast, the general notion is supported both by biophysical considerations^21^ and by imaging cells along the long axis, where they appear round^22-24^. Furthermore, Pomegranate’s approach requires that the long axis of cells is parallel to the XY imaging plane. This will typically be the case when cells are imaged in microfluidics channels, but the requirement may not always be met when cells are fixed on glass slides without any constraint in Z.

Pomegranate uses full voxels rather than sub-voxel segmentation for ROIs (Fig. 2B). Calculation of volume and surface area could be refined by using the exact rather than voxelized shape, but this calculation has not been implemented. Furthermore, Pomegranate is currently only compatible with single timepoint images. Using the pipeline for time series images will require analyzing each timepoint individually and linking the resulting ROIs.

## Summary

Because Pomegranate can work with both externally and internally generated 2D segmentations, we see it not as an alternative but as a valuable extension to the image analysis portfolio that many groups already use. Pomegranate’s 3D reconstruction algorithm improves volume estimates and allows for quantification of cellular signals in 3D (Fig. 6). Since the only assumption for the 3D extrusion is radial symmetry around a long axis, Pomegranate is applicable to all cell types with such shape including many rod-shaped bacteria in addition to fission yeast.

## Code availability

The software, usage guide, example images, and additional tools are available at the Pomegranate GitHub repository (https://github.com/erodb/Pomegranate). Installation of Pomegranate is possible through a FIJI/ImageJ update site (https://sites.imagej.net/HaufLab) or through direct download of the macro files from GitHub.

## Materials and methods

### Fission yeast strains

| Strain Name | Mating Type | Genotype | Used in |
| --- | --- | --- | --- |
| SW184 | *h-* | *leu1 Z<<natR<<Padh31-tetR-tdTomato* | Fig. 4, 5, S2, S3, S4, S5, S6, S7, S8 |
| SW169 | *h+* | *Z<<natR<<Padh31-tetR-tdTomato mep33+-mCherry<<hygR* | Fig. 3, S5 |
| SW191 | *h+* | *Z<<natR<<Padh31-tetR-tdTomato mep33+-tdTomato<<kanR* | Fig. 3, S5 |
| SW178 | *h+* | *mad1+-ymEGFP Z<<natR<<Padh31-tetR-tdTomato* | Fig. 4, S2, S6, S7, S8 |
| SW179 | *h+* | *mad1-codonOpt-ymEGFP Z<<natR<<Padh31-tetR-tdTomato* | Fig. 4, S3, S6, S7, S8 |
| SW180 | *h+* | *mad2+-ymEGFP Z<<natR<<Padh31-tetR-tdTomato* | Fig. 4, S2, S4, S6, S7, S8 |
| SW181 | *h+* | *mad2-codonOpt-ymEGFP Z<<natR<<Padh31-tetR-tdTomato* | Fig. 4, 5, S2, S4, S6, S7, S8 |
| SW182 | *h+* | *mad3+-ymEGFP Z<<natR<<Padh31-tetR-tdTomato* | Fig. 4, S6, S7, S8 |
| SW183 | *h+* | *mad3-codonOpt-ymEGFP Z<<natR<<Padh31-tetR-tdTomato* | Fig. 4, S2, S3, S6, S7, S8 |
| JX798 | *h-* | *ade6-M210 ura4-D18 tea1Δ::ura4+* | Fig. 5, S9 |
| JY327 | *h-* | *wee1-50* | Fig. 5, S9 |
| SW323 | *h+* | *cut1+-GFP<<kanR cdc25-22* | Fig. 5, S9 |

The *tea1Δ::ura+^57^, wee1-50^58^, cdc25-22^59^ and tetR-tdTomato-NLS^39^* alleles have been described, and strains expressing these were obtained from Y. Watanabe. The GFP-tags present in some strains are not relevant for the purpose of this study. Yeast monomeric enhanced GFP (ymEGFP) tagged alleles of *mad1, mad2 and mad3* will be described elsewhere (Esposito et al., in preparation).

### Microscopy

Fission yeast cells were revived from frozen stocks on YEA plates (yeast extract with additional 0.15 g/L adenine)^67^. For imaging, cells were inoculated into EMM2 (Edinburgh Minimal Medium, MP Biomedicals # 4110012)^67^ with 0.2 g/L leucine or 0.15 g/L adenine added for auxotrophic strains. Cells were grown to a concentration of around 1 × 10^7^ cells/mL. Cells were diluted to a concentration of 3 × 10^6^ cells/mL and loaded into the cell input wells of a CellASIC ONIX Y04C microfluidics plate. EMM2, with leucine or adenine if needed, was loaded into the media input wells of the microfluidics plate.

Cells were loaded into the imaging window of the microfluidics plate by multiple 5 second pulses of 8 psi. The six media input wells of the microfluidics plate were run in symmetric pairs (1/6, 2/5, and 3/4) to maintain homogenous flow direction. Medium was run at 6 psi.

The microfluidics plate was imaged using an inverted widefield DeltaVision Microscope with pco.edge 4.2 (sCMOS) camera, Olympus 60X/1.42 (UIS2, 1-U2B933) objective, Trulight fluorescent illumination module, and EMBL environmental chamber. Imaging was performed at 30°C (34°C for fission yeast *cdc25-22* mutants).

Multi-channel Z-stack images were collected with 72 or 100 optical sections, at a section spacing of 0.1 μm. With these parameters, a 0.1071 μm (X axis) x 0.1071 μm (Y axis) x 0.1 μm (Z axis) voxel size is generated. Note that axial distances are recalibrated later (Fig. S1, S2). Images were taken at multiple locations within the microfluidics chamber, sampling across a wide range of cell densities.

### Image corrections and calibrations

All raw images were deconvolved with three cycles of conservative deconvolution, without additional deconvolution corrections or normalizations, using softWoRx (Applied Precision). Deconvolved images were subjected to flat-field correction, axial chromatic aberration correction (Fig. S1A,B), and axial distance correction (Fig. S1C-F).

Flat-field correction was applied to correct for non-uniform illumination of the imaging field. Images for flat-fielding were acquired by imaging Alexa Fluor-488 and Alexa Fluor-568 labeled antibodies (Invitrogen/Thermo Fisher) diluted 1:100 in PBS and mounted in an 8-well μ-Slide (Ibidi). Flat-field images from different sample positions were averaged to compensate for stochastic differences. Dark noise images were collected by imaging with a closed shutter. Dark noise was subtracted from all flat-field and raw images. Flat-field images were normalized by dividing all pixel intensities by the mean image intensity. Finally, dark noise-subtracted raw images were divided by dark noise-subtracted, normalized flat-field images.

In order to correct for axial chromatic aberration (i.e. light of different wavelengths getting focused onto different optical sections), 0.1 μm TetraSpeck Fluorescent Microspheres (Invitrogen/Thermo Fisher) were imaged (Fig. S1A). The beads were viewed from the XZ orthogonal perspective, and the chromatic aberration along the Z axis was quantified in Fiji by manually measuring the distance in pixels between the centers of the bead image in each fluorescent channel (Fig. S1B). Raw images were corrected by shifting the Z position of the optical sections for each channel relative to the other channels based on the measured axial chromatic aberration.

We noted that our images had an axial distortion that made round objects appear elliptic in XZ or YZ projections (Fig. S1C,E), presumably a consequence of using a widefield system. In order to correct for this axial distortion, 4 μm TetraSpeck Fluorescent Microspheres were imaged in the same medium that was used for cells (EMM2). The beads were viewed from the XZ orthogonal perspective, and the geometric elongation was quantified manually in Fiji by measuring the width and height of the bead (Fig. S1D). The ratio of these was used as the correction factor. Raw images were corrected by multiplying the Z distance between planes with the correction factor to undo the apparent stretching in Z. Other imaging systems may not require such a correction. When we used TetraSpeck Fluorescent Microspheres pre-mounted on a slide to measure the correction factor, the value was higher (Fig. S1F) and insufficient to lead to a round appearance of nuclei in XZ or YZ. If such a correction is needed, we therefore recommend imaging beads in the same medium as cells, or empirically determining the factor using the assumption that nuclei have a round shape in orthogonal views.

Because the strategies for nuclear and whole-cell 3D segmentation are different, nuclear and whole-cell volumes are differently affected by the correction factor (Fig. S2B). The volume of nuclei measured in voxels stays constant but nuclear volume in μm^3^ changes. In contrast, because cell volume is constructed by extrusion, the cell volume in μm^3^ stays constant, regardless of the correction factor, but the number of voxels in the 3D ROI changes.

### Pipeline development and testing

Pomegranate was written in the macro language of the ImageJ/Fiji open-source imaging processing software^3^. Development and testing was performed on a 2017 iMac (4.2 GHz Intel Core i7 Processor, 16 GB 2400 MHz DDR4 RAM, Catalina Version 10.15.5) and a 2018 Mac mini (3 GHz Intel Core i5, 16 GB 2667 MHz DDR4 RAM, macOS Mojave Version 10.14.6).

### Nuclear 2D segmentation by Pomegranate

Nuclear segmentation was accomplished by using a fluorescent marker enriched in the nucleus (TetR-tdTomato-NLS^39^). Images were taken at multiple optical sections and segmentation was performed on each section. A Gaussian blur and unsharp mask were used to enhance acutance and edge contrast for the nuclei (Fig. 1A, panel 2). An adaptive threshold (Otsu’s method^68^) was used to produce a binary image (Fig. 1A, panel 3). A Gaussian blur and remasking was used to smooth the binary image (Fig. 1A, panel 4). Fiji’s “Analyze Particles” algorithm was used to translate the binary image into nuclear ROIs.

To eliminate 2D ROIs that deviate too much from the expected idealized circular shape, we calculated circularity, *C* as

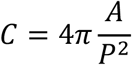

where A is the ROI area, and P is the ROI perimeter. ROIs below a circularity of 0.6 were excluded by Fiji’s “Analyze Particles” algorithm. ROIs with a low circularity typically occurred when the nucleolar signal was too weak and had not been incorporated into the segmentation, resulting in a crescent-shaped region. We further dilated all nuclear ROIs by 0.1 μm, which was necessary to improve acquisition of target fluorophores at the nuclear periphery. Dilation is optional in the pipeline and the extent of dilation can be defined by the user. Finally, an elliptical fit was performed on all nuclear ROIs, to alleviate the impact of irregular shapes resulting from lower nucleolar signals. Elliptical fits convert the ROI into an ellipse where the area, orientation and centroid of the original ROI are preserved.

### Nuclear 3D reconstruction by Pomegranate

For nuclear reconstruction, ROIs from the nuclear 2D segmentation were grouped by their centroid coordinates to form nuclei (Fig. 1A, panel 5). ROIs in adjacent optical sections were grouped when the XY distance between their centroids did not exceed 2.0 μm. To eliminate nuclei that had moved during acquisition, nuclei were excluded if the XY distance of any single centroid to the mean nuclear centroid exceeded 3.0 μm. These threshold values can be changed by the user. In instances where nuclei had intermittent optical sections without ROI, the largest subset of contiguous nuclear ROIs was isolated. Nuclei were removed when the total number of 2D ROIs was less than 5. For each nucleus, the ROI with the largest area was designated the nuclear midplane. Nuclei were excluded if this ROI was at the top or bottom of the ROI stack. Nuclear volumes were calculated as the sum of the volumes of nuclear ROI voxels (using the Z thickness obtained after correction for axial distortion).

### Whole-cell 2D segmentation by Pomegranate

When a bright-field image is provided, Pomegranate uses its default 2D segmentation algorithm to produce the 2D binary required for 3D reconstruction. Initial whole-cell segmentation was accomplished by first reducing the brightfield image Z-stack to the optical sections below the most in-focus optical section, which we defined as the optical section for which the standard deviation of pixel intensities in the image is minimized (Fig. S3). Since we used a microfluidics chamber in which cells form a flat layer, we initially assumed that the most in-focus optical section is the same for all cells in an image. A Gaussian blur was used on each optical section to reduce image noise. An unsharp mask was used to enhance the acutance of the boundary band surrounding each cell. An adaptive threshold (Otsu’s method^68^) was used to produce a binary image for each optical section. In optical sections close to the focal plane, cell boundaries are hard to detect as the boundary band width approaches zero (Fig. S3A, 0 μm). The farther an optical section is from the focal plane, the larger the boundary band width becomes (Fig. S3A, −2.4 μm). We therefore averaged the binary images from all remaining optical sections to produce an image where boundary bands appear as gradients, akin to valleys in a topographic map (Fig. 1B, panel 2). We interpret the darker regions in this grayscale map to be high confidence areas for cell borders. An additional adaptive threshold was used on this projection (Otsu’s method^68^). Using these steps, cells were surrounded by a boundary band (Fig. 1B, panel 3). Fiji’s “Analyze Particle” tool was used to translate the binary image into whole-cell ROIs, and a distance transform was used to measure the thickness of the boundary bands. All ROIs were then enlarged by the half-width of the boundary band (Fig. 1B, panel 4). An optional watershedding algorithm (Watershed Irregular Features, BioVoxxel Toolbox^69^) in the pipeline allows for separating unsegmented adjacent cells or regions.

Debris or cell features such as septa and vacuoles can introduce gaps or jagged edges in the segmented brightfield binary. To compensate for this, a smoothing algorithm was used. The smoothing algorithm closes gaps in the ROI that fall beneath a size threshold defined by the user. Additionally, cells along the image periphery were excluded based on a user-defined margin.

Adapted from the MAARs pipeline^33^, an optional solidity filter was used to exclude some ROIs that do not represent cells. An ROI’s solidity is the ratio of its area and its convex hull area, where a value of 1 indicates a completely convex object. We made this step optional and adjustable, because, as also observed by Liu *et al*.^34^, cells with curvature may be falsely excluded while intercellular regions with high solidity may be falsely included. An additional manual exclusion step allows users to remove falsely included regions prior to 3D reconstruction.

Within the pipeline, whole-cell 2D ROIs are initially constructed in the optical section that is considered the most in-focus for the entire image. However, this does not always coincide with the optical section where the nucleus has its largest diameter (which we consider the midplane for the nucleus). Differences between the nuclear and cellular midplane position result in vertical misalignment between a nucleus and its enclosing cell. As an additional optional step, when nuclear data are available, the whole-cell 2D ROI for a given cell can therefore be re-assigned to the midplane of the corresponding nucleus, which we did when nuclear measurements were available (Fig. 4, but not Fig. 5).

### Whole-cell 2D segmentation by YeaZ

To provide an alternative 2D segmentation input for Pomegranate whole-cell reconstruction, the YeaZ neural-network-driven segmentation pipeline was used^41^. An in-focus slice was isolated from the brightfield Z-stack and used as input for the YeaZ pipeline. Threshold value was set to 0.5 and the minimum distance between seeds was set to 10. The binary image was further refined with watershedding using a separator size of 20 pixels (Watershed Irregular Features, BioVoxxel Toolbox^69^). The resulting 2D binary image was used as input for Pomegranate whole-cell reconstruction.

### Whole-cell 3D reconstruction by Pomegranate

We used the assumption that cells are radially symmetric around a central axis along the length of the cell^21^ to guide our whole-cell 3D reconstruction. To reconstruct cells, we use the midplane ROI as the starting point. The reconstruction algorithm then extrudes a series of spheres along the topological skeleton of the cell (Fig. 2A).

The Euclidean distance map and the topological skeleton were generated from the binary midplane ROI. The Euclidean distance map is a greyscale image where the value of each pixel is its Euclidean distance from the closest edge of the ROI (Fig. 1B, panel 6; Fig. 2C). We used the Fiji implementation of an algorithm for generating a Euclidean distance map^70^. The topological skeleton is a one-pixel-wide shape that is equidistant from the boundaries of the ROI (Fig. 1B, panel 5). We used the Fiji legacy implementation of a thinning algorithm^71^ to produce topological skeletons. The union of the distance map and topological skeleton yields a topological skeleton where each pixel along the skeleton has a value representing the distance of the topological skeleton to the ROI boundary (Fig. 1B, panel 7; Fig. 2C). For 3D extrusion (Fig. 2A), we centered spheres on these pixels with the radius of the sphere taken from the distance value of each pixel (Fig. 2C,D). Due to a limitation in Fiji’s ROI system, ROIs are always two-dimensional, confined to a single optical section. In order to construct a sphere, concentric, circular, cross-sectional ROIs of the sphere are created whose radii are a function of the parent sphere’s radius, and the cross section’s axial distance from the midplane (Fig. 2B). The cross-sectional radii are given by

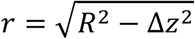

where *r* is the radius of the cross-section ROI, *R* is the radius of the parent sphere, and *Δz* is the cross-section’s distance from the midplane. Sub-pixel coordinates for these cross-sectional ROIs were resolved by rounding to their nearest pixel coordinates (Fig. 2B). The union of all coplanar circular cross sections forms the 2D ROI for a given optical section. All 2D ROIs for a given cell were associated across optical sections to form the whole-cell 3D ROI (Fig. 2E). The amount of detail in the reconstruction is proportional to the resolution of the imaging system (Fig. 2F).

### Whole-cell and nuclear reconstruction pairing

Whole-cell and nuclear reconstructions, if processed simultaneously in Pomegranate, can be paired on a cell-by-cell level. To accomplish this, Pomegranate extracts the centroid of each nucleus, and identifies the whole-cell ROI that contains that centroid. Only whole-cell ROIs paired with a single nuclear centroid are kept. Cellular ROIs with two or more nuclei (septating cells or poor segmentation), or cells without nucleus (background ROI or failed nuclear segmentation), are excluded. Nuclei without an associated whole-cell ROI are also excluded. Optionally, we aligned nuclear and whole-cell reconstruction by assigning the nuclear centroid Z position as midplane of the whole-cell ROI.

### Calculation of volumes and surface areas from Pomegranate ROIs

For both nuclear and whole-cell 3D analyses, volumes were calculated as the sum of the volumes of the ROIs in each optical section (area of the 2D ROI, multiplied by the Z thickness, using the Z thickness obtained after correction for axial distortion).

For both nuclear and whole-cell analyses, surface area *S* is given by

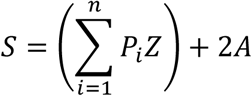

Where is *P*_*i*_ is the perimeter of an ROI, for *n* ROIs in a single cell, *A* is the area of the midplane ROI, and *Z* is the spacing between optical sections along the Z-axis. 2*A* is equivalent to the area of all surfaces in the reconstruction parallel to the XY plane, whereas the sum of all *P*_*i*_*Z* is equivalent to the area of all surfaces orthogonal to the XY plane.

### Calculation of whole-cell volumes and surface areas from idealized rod shape

To calculate shape descriptors for idealized, rod-shaped cells (Fig. 4A), we extracted cell length and width for these idealized cells from the midplane ROI using two different methods. The first method used the major and minor axes of an elliptical fit as cell length (*L*) and width (*W)*. The second method used the maximum and minimum Feret diameter (also called the caliper diameter) as length and width. The Feret/caliper diameter is the distance between two parallel lines tangential to the ROI. Cells were assumed to be rod-shaped, described as a cylinder with hemispheres at either end. The volume is given by:

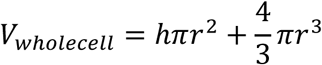

where *h* is the length of the cylindrical body and *r* is the radius of the cell from the central axis. Radius and length of the cylindrical body were calculated from cell width and length as *r = W / 2 and h = L – 2r*.

The surface areas of whole-cell reconstructions are given as the sum of the surface areas of the lateral face of a cylinder and of the hemispheric ends:

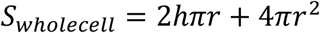

where *h* is the length of the cylindrical body and *r* is the radius of the cell from the central axis. Radius and length of the cylindrical body were calculated from cell width and length as *r = W / 2 and h = L – 2r*.

### Calculation of nuclear volumes and surface areas from idealized elliptic shape

For nuclei, the longer (*L*) and shorter (*W*) diameter were determined by elliptical fit or Feret diameters, similar to the parameter extraction described for whole-cell idealized volumes. Volumes were calculated under the assumption of nuclei being ellipsoidal. The volume is given by:

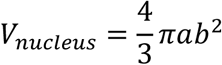

with *a* = *L* / 2 and *b* = *W* / 2.

The surface areas of the same nuclei are given by the following approximation for the surface area of an ellipsoid:

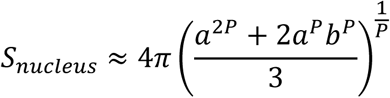

where *a = L / 2* and *b = W / 2*, and *P* = 1.6075.

### Calculation of nuclear and whole-cell fluorescence intensities

To quantify fluorescent signal within ROIs, the 2D cross-sectional ROIs are used to measure the integrated density of each region. The integrated density across all cross-sections is summed up to produce the volumetric integrated density. To calculate mean intensity in each region, volumetric integrated density is divided by the volume of the 3D ROI.

### Downstream processing

Compilation, processing, and data visualization of Pomegranate outputs were performed in R^72^, using the tidyverse^73^, ggpubr^74^, ggcorrplot^75^, MASS^76^ and gridExtra^77^ packages.

For any Pomegranate analysis reporting whole-cell as well as nuclear data, we only included cells with both nuclear and cellular segmentation (Fig. S2A). For the analysis of cells with wild-type morphology (Fig. 4, Fig. S6-8), we applied an additional, optional filter that only includes cells when both cell and nuclear 3D ROI have an aspect ratio between 0.8 and 1.2 (Fig. S2A). We define the aspect ratio as extension of the region in Z divided by the largest diameter or cell width in X/Y. Aspect ratios of less than 1 occur when a cell is shifted in Z and the collected optical sections do not fully cover the cell volume (Fig. S2A). Imperfect nuclear acquisition can result in aspect ratios higher than 1.2 (Fig. S2A). In these cases, the nuclear midplane is often asymmetrically located, which may misplace the whole-cell segmentation with respect to the nucleus. This filter was not applied to datasets where reconstructions were manually inspected (Fig. 3, Fig. S5), or to datasets of cell shape and size mutants (Fig. 5, Fig. S9).

For the analysis of cells with wild-type morphology (Fig. 4, Fig. S6-8), we further excluded cells whose nuclear ROI extended beyond the boundaries of their associated whole-cell ROI. This filter could not be applied to the other analyses, where only whole-cell or only nuclear data was collected.

## Acknowledgements

We thank the Nurse and Watanabe labs, as well as F. Obuekwe and D. Weidemann for generating yeast strains, A. Brock and B. Jutras for providing images of bacteria, D. Weidemann for feedback on the pipeline, J. Rogers and C. Morton for assisting in experiments related to performance testing, E. Wood for recommendations on microfluidics settings, Wayne Rasband and the Image.sc Forum community for their guidance and support, and all members of the Hauf lab for comments on the manuscript.

## Author Contributions

Project administration and supervision: SH. Experiment design: EKB, EE, SH. Imaging: EKB, EE. Pipeline programming: EKB. Pipeline testing and evaluation: EKB, EE. Pipeline documentation: EKB. Manuscript writing: EKB, SH. Manuscript revision: EKB, EE, SH.

## Additional Information

The authors declare no competing interests.

