## Supplemental Figures S1 - S10 for "Pomegranate: 2D segmentation and 3D reconstruction for fission yeast and other radially symmetric cells"

### Baybay Figure S1

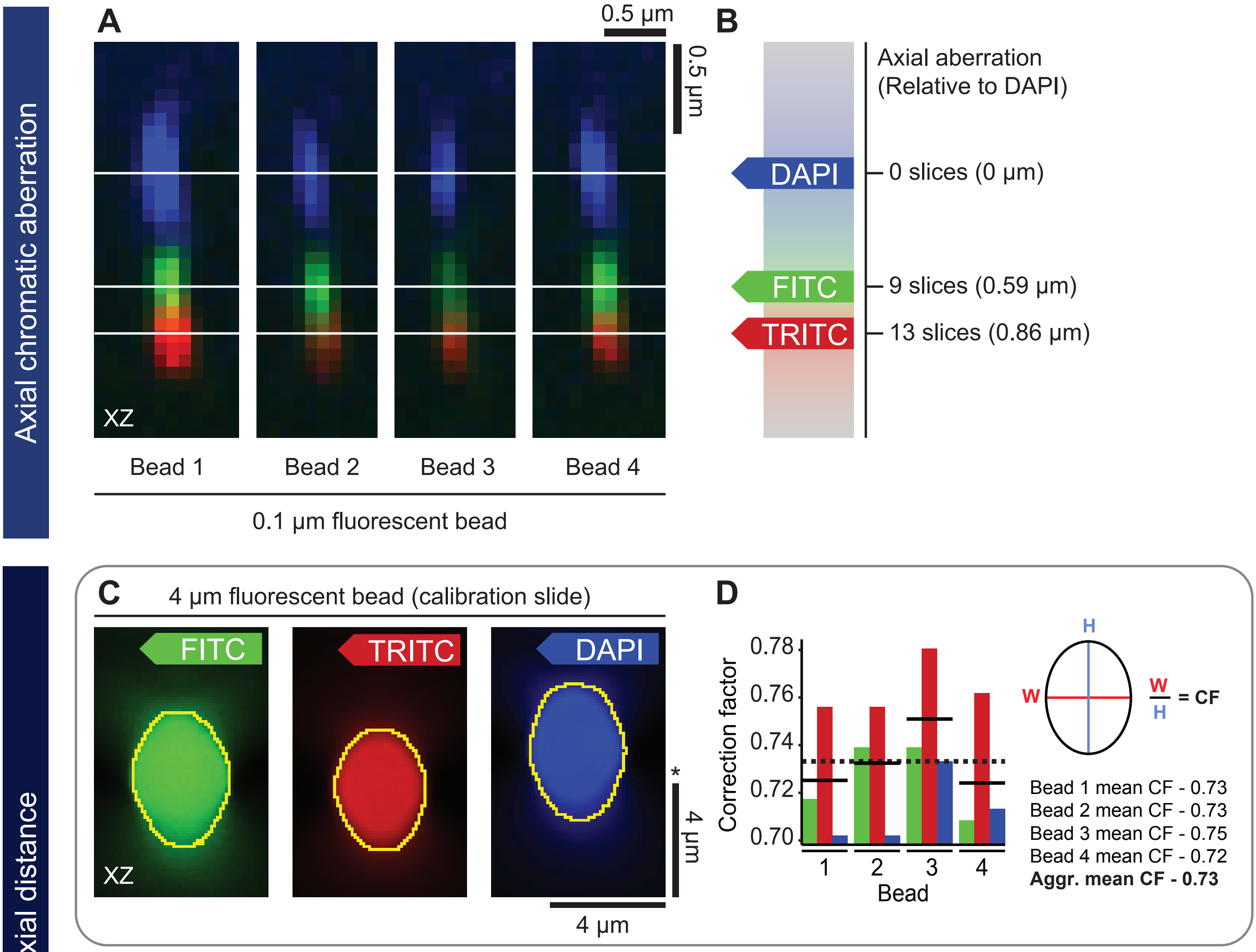


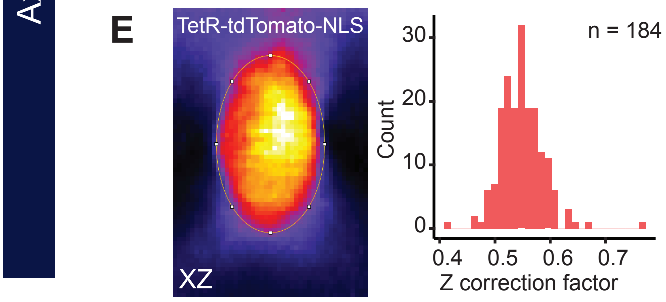


**F**

| Method | Z correction factor |
|---|---|
| 4 µm beads (EMM2) | 0.66 |
| 4 µm beads (calibration slide) | 0.73 |
| Nuclei spheroid calibration | 0.58 |

**Supplemental Figure S1. Axial corrections** - Both axial chromatic aberration and distances need to be corrected for in order to obtain accurate quantifications of fluorescence intensities. **(A)** Correcting for axial chromatic aberration: XZ maximum projections of 0.1 µm TetraSpeck Fluorescent Microspheres. White bars are manually annotated midslices for each channel. Note that the vertical scale bar has been corrected with the appropriate axial distances (described in C-F). **(B)** Quantification of axial aberration relative to DAPI (blue fluorescence). **(C)** Sample axial distance calibration approach using fluorescent 4 µm beads. Shown are uncorrected XZ maximum projections of 4 µm TetraSpeck Fluorescent Microspheres. Note that the vertical scale bar (asterisk) represents uncorrected axial distance. **(D)** Bar plot of correction factors (CF) calculated as the aspect ratio (W/H, width/height) of the beads in XZ across multiple channels. Solid lines indicate mean CF for each bead, dashed line indicates mean CF across all beads (n = 4). **(E)** Alternative approach to axial distance calibration leveraging the assumption that nuclei are spherical in wild-type fission yeast: XZ maximum projection of a single fission yeast nucleus with TetR-tdTomato-NLS without correcting axial distance (left) and a histogram (bins = 30) of required CFs to yield spherical nuclei, from an image with 184 nuclei. **(F)** Table of Z correction factors derived from the methods described in C-E.

### Baybay Figure S2

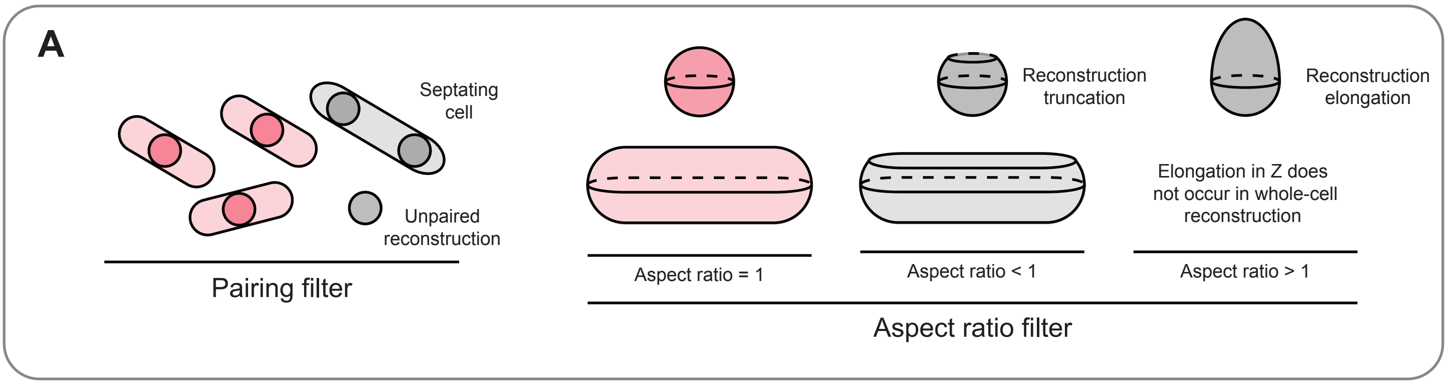


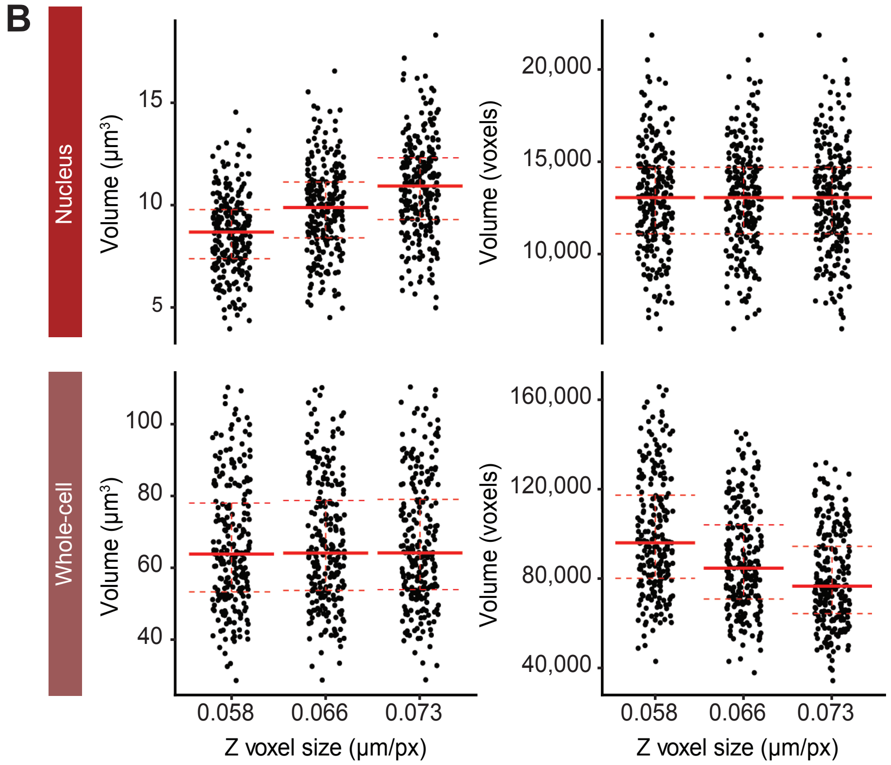


**C**

| Z | Volume | Nucleus median | Nucleus SD | Whole-cell median | Whole-cell SD |
|---|---|---|---|---|---|
| 0.058 | µm3 | 8.7 | 1.9 | 63.8 | 17.4 |
| | voxels | 13,056 | 2,858 | 95,960 | 26,117 |
| 0.066 | µm3 | 9.9 | 2.2 | 64.1 | 17.5 |
| | voxels | 13,056 | 2,858 | 84,673 | 23,084 |
| 0.073 | µm3 | 10.9 | 2.4 | 64.2 | 17.5 |
| | voxels | 13,056 | 2,858 | 76,622 | 20,896 |

**Supplemental Figure S2. Comparison of multiple Z voxel sizes - (A)** Cartoon depicting the two common filters applied to Pomegranate outputs: pairing and aspect ratio. The pairing filter lets only objects pass that contain one nuclear ROI and one whole-cell ROI. Aspect ratio is calculated as the ratio of height in the Z direction against width in the XY plane. A low (<1) aspect ratio typically occurs when the midplane is not centered and not enough optical sections are available, so that the reconstruction is truncated. A high (>1) aspect ratio for nuclear ROIs typically occurs when background flouresence is mistakenly included. High aspect ratios do not occur for whole-cell ROIs due to the radial symmetry of the spherical extrusion. When applying the aspect ratio filter, we included ROIs with aspect ratio between 0.8 and 1.2. **(B)** Plots of whole-cell and nuclear reconstruction volumes (both in cubic microns and in voxels) for identical sets of cells, processed in Pomegranate with different Z voxel sizes (determined from the analyses in S1) with data filtered as described in (A). Each point represents one cell. Overlaid lines represent median (solid red line), as well as the first and third quantile (bottom and top dashed red lines, respectively). Because of the different modes of 3D reconstruction, nuclear and cellular ROI voxel numbers and volumes in cubic microns are differently affected by the correction factors. **(C)** Table of descriptive statistics from (B) (SD = standard deviation).

### Baybay Figure S3

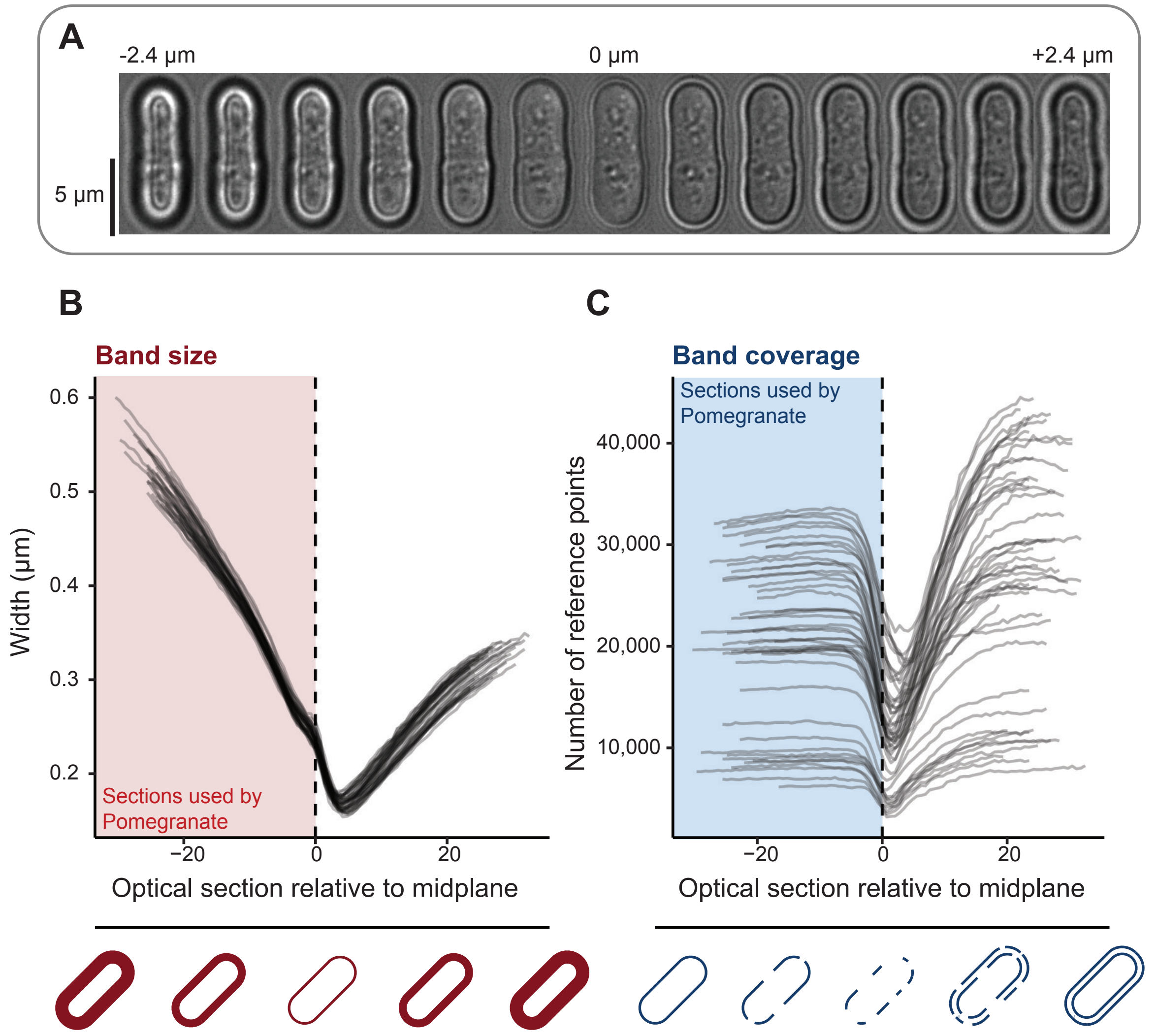


**Supplemental Figure S3. Features of the brightfield image Z-stack - (A)** Sample brightfield images at various optical sections in Z. **(B)** Line plot of mean black band width in a full field as a function of Z position, for multiple images. Shaded regions indicate the opticals sections used in Pomegranate's default 2D segmentation. The dashed line indicates the optical section in which the standard deviation of brightfield intensities is minimized – which we interpret as the most in-focus optical section. **(C)** Line plot of number of reference points used in black band width measurement in a full image as a function of Z position. The number of reference points is the number of pixels in the topological skeleton of a binarized black band (by adaptive threshold) – and is a metric of the overall length of the black band. The black band disappears as the brightfield image comes into focus. Dashed vertical line indicates the optical section in which the standard deviation of intensities is minimized. Schematic drawings at the bottom illustrate the reason for the changes in y-axis value.

### Baybay Figure S4

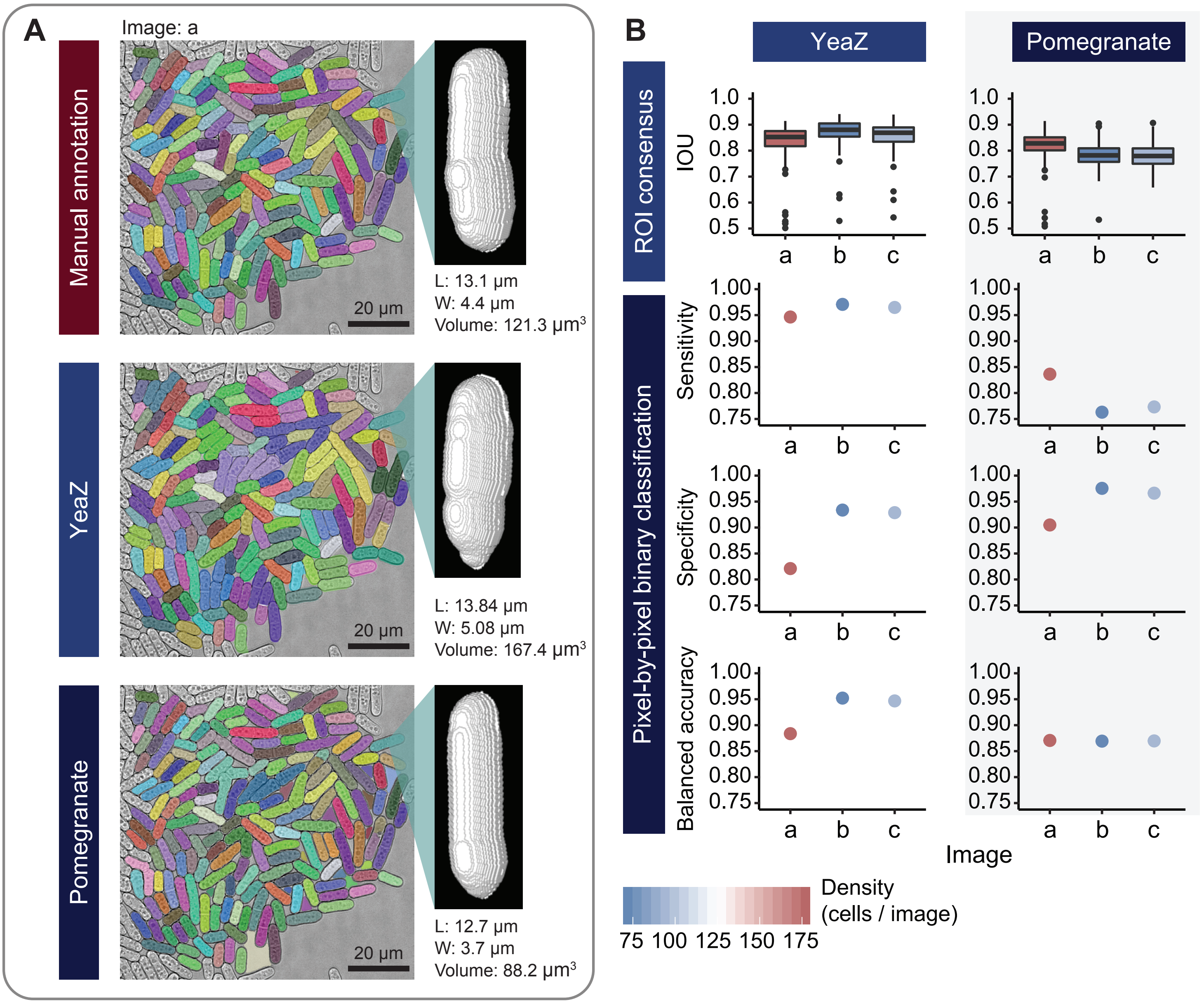


**Supplemental Figure S4. Comparison of 2D segmentation approaches - (A)** Sample brightfield image with overlaid ROIs generated by manual annotation (ground truth), YeaZ neural network-based segmentation, and Pomegranate's default 2D segmentation algorithm. Cell colors are conserved across images. YeaZ or Pomegranate ROIs are considered to be targeting the same ground-truth cell if the intersection-over-union (IOU) with the ground-truth ROI surpasses a value of 0.5. YeaZ or Pomegranate ROIs unpaired with a ground-truth ROI were assigned a random color. Images shown represent image "a" from the three images (a, b, c) analyzed in (B). Each image is accompanied by a sample 3D reconstruction of the same cell using the ROIs generated by each of the three methods. Annotations provide size parameters for each cell: maximum Feret diameter (length, L) minimum Feret diameter (width, W), and whole-cell volume. **(B)** Quantification of 2D segmentation performance across three images using two tests: ROI consensus and pixel-by-pixel binary classification. Image "a" is shown in (A). Boxplots illustrates IOU of Pomegranate or YeaZ ROIs compared to their corresponding ground-truth ROI. Boxes represent inter-quartile range (IQR), while whiskers extend to the smallest and largest value no further than 1.5 times the IQR from the first and third quartile, respectively. Points beyond the whiskers are registered as outliers and plotted individually. Horizontal bars at the center of boxes indicate medians. Pixel-by-pixel binary classification measures the ability for a segmentation approach to correctly classify individual pixels between being part of a cell (positive), or part of the intercellular space (negative). Cells that intersect a 10 pixel margin with the edge of the image were excluded from the analysis (grey in A). Sensitivity was calculated as the number of pixels correctly classified as cellular (true positives) divided by all pixels that are cellular in the ground truth manual annotation (true positives + false negatives); specificity was calculated as the number of pixels correctly classified as non-cellular (true negatives) divided by all pixels that are non-cellular in the ground truth manual annotation (true negatives + false positives). Balanced accuracy is the mean of sensitivity and specificity. Color of boxes or points represents image density (cells/image).

#### Baybay Figure S5

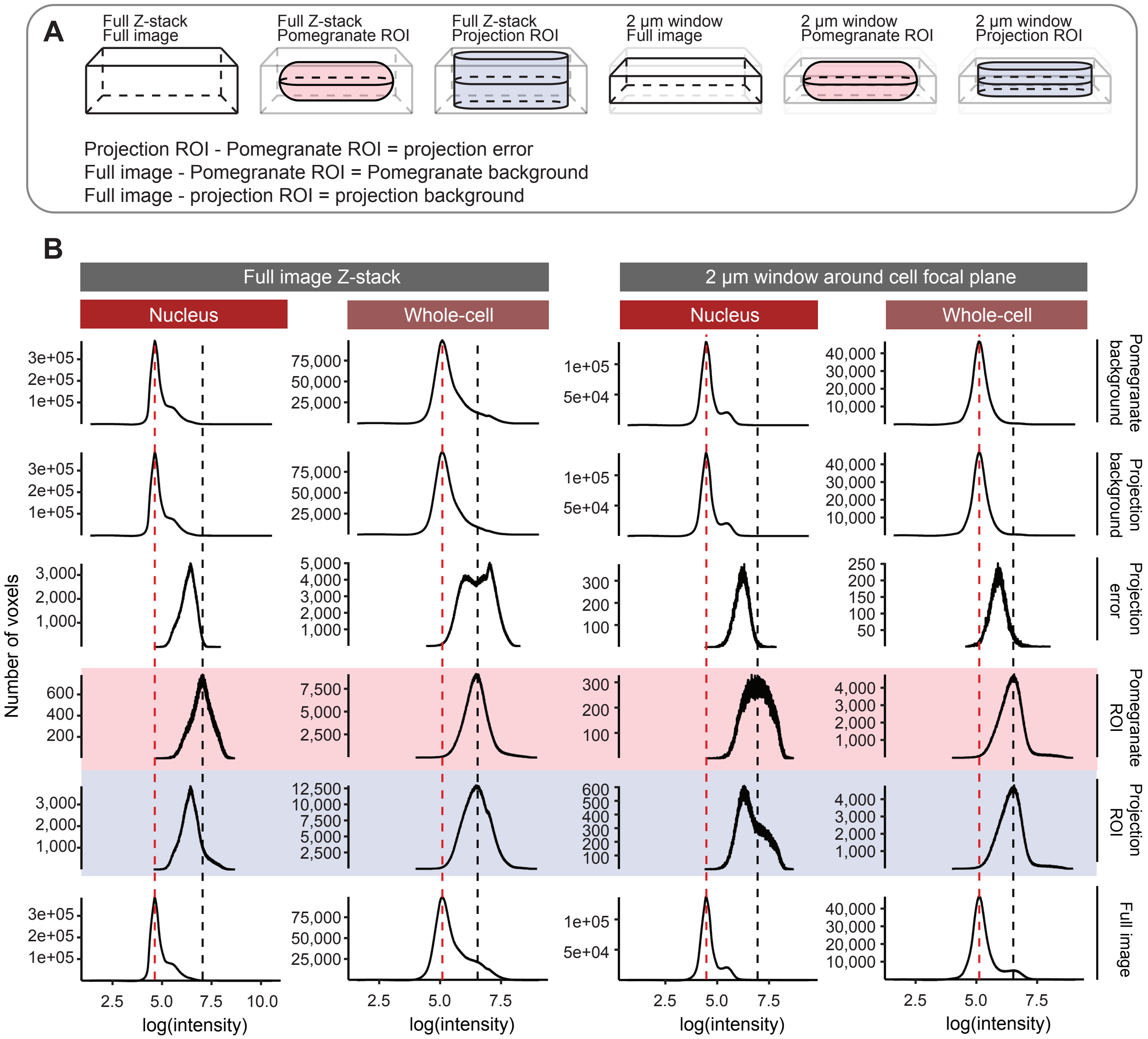


**Supplemental Figure S5. Detailed signal acquisition performance - (A)** Cartoon representing regions described in the rows of (B). 'Pomegranate ROI' refers to the output of Pomegranate's reconstruction algorithm. 'Projection ROI' refers to the region where the midplane ROI is used as ROI in all Z-sections. Projection error is the subtraction of signal acquired from Pomegranate's ROI from the signal acquired by the projection method. 'Pomegranate background' and 'Projection background' are the voxels in the full image not contained within either the Pomegranate ROI or the Projection ROI, respectively. Note that full images contain multiple cells. **(B)** Intensity histograms as in Figure 3B, with additional details. Either the full image Z-stack is used for analysis (left), or analysis is restricted to a 2 µm window in Z around the midplane (right). Dashed lines annotate maxima: background (red) and signal of interest (black). Use of projection collects lower intensities (likely extra-nuclear or extra-cellular) signals. Higher intensity signal acquired by the projection approach (seen as the second, higher-intensity maximum in the projection error in the full Z-stack) is likely caused by unbleached autofluorescence, which tends to disappear after a few exposures.

### Baybay Figure S6

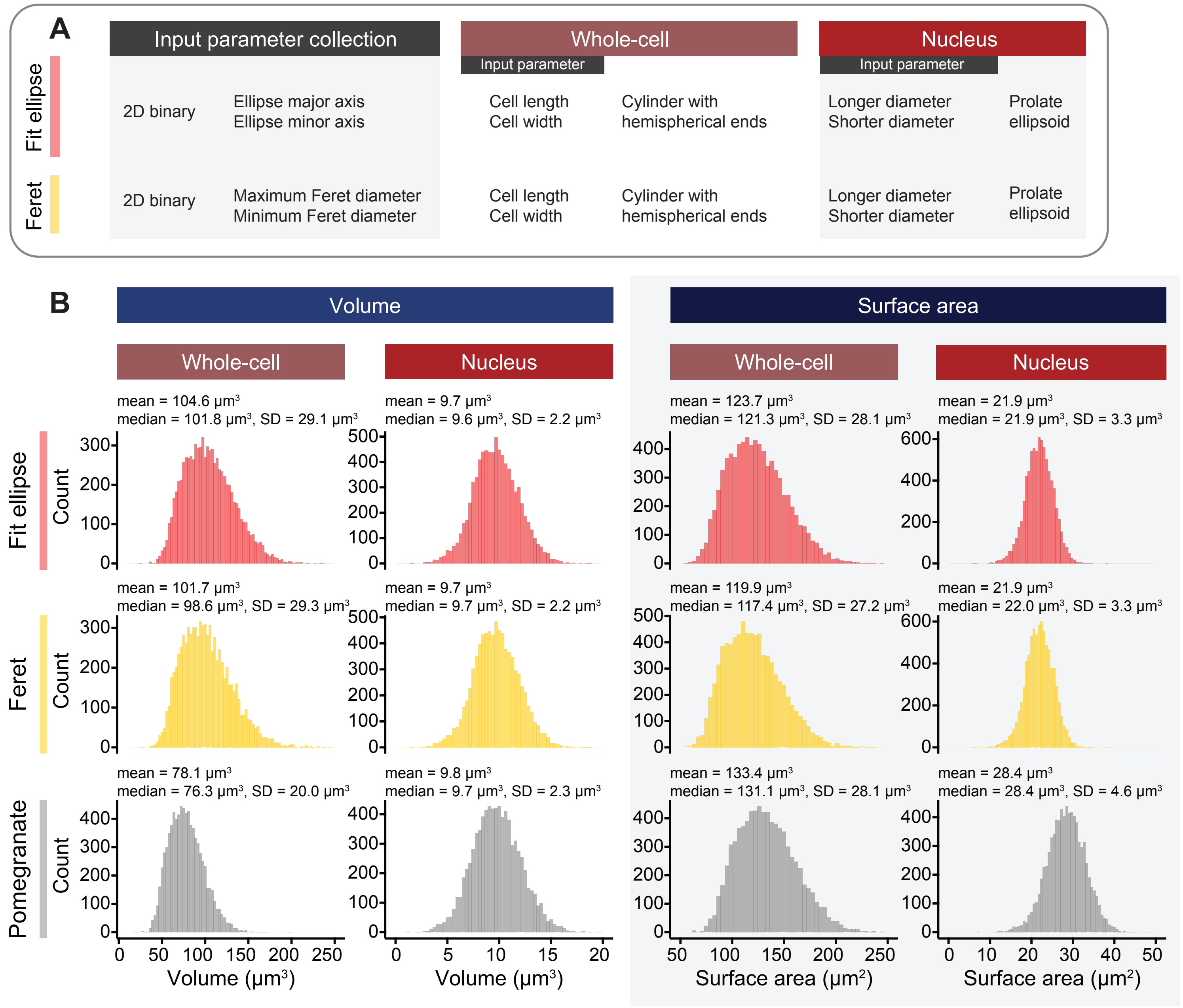


**Supplemental Figure S6. Analysis of volumes and surface areas - (A)** Tabular schematic of idealized volume reconstruction approaches. For whole-cell idealized volumes, cell length and cell width are inputs for a rod-shaped geometry. Cell length and width are extracted from the 2D midplane ROI via the major and minor axes of a fit ellipse, or via the maximum and minimum Feret diameter, respectively. For nuclear idealized volumes, the longer and shorter diameter for the idealized nucleus are collected in the same way as for whole-cell ROIs using an elliptical fit or the Feret diameters. Longer and shorter diameter are inputs for a prolate ellipsoid with the longer diameter parallel to the cell midplane. **(B)** Histograms of whole-cell and nuclear volumes and surface areas of wild-type cells (n = 8,026), comparing multiple strategies for reconstructing fission yeast volume. Cells were included based on the filters described in Figure S2A, in addition to removing cells where the nucleus extended beyond the border of the cell. Whole-cell volume column binwidth = 3.0; nuclear volume column binwidth = 0.36; whole-cell surface area column binwidth = 4.0; nuclear surface area column binwidth = 0.68 (Freedman-Diaconis). Annotations provide descriptive statistics (SD = standard deviation). Volumes and surface areas for idealized volumes are calculated as a cylinder with hemispherical ends (cells) and prolate ellipsoid (nuclei). Pomegranate volumes and surface areas are derived from the Pomegranate reconstruction algorithm. Pomegranate volumes are given by the sum of the cross-sectional areas of all optical sections in a cell, multiplied by the spacing between optical sections in Z. Pomegranate surface area is given by the sum of perimeters of all optical sections in a cell, multiplied by the spacing between sections (lateral surface area) - added to the sum of all exposed top and bottom faces in the reconstruction - which is equal to two times the midplane cross-sectional area.

### Baybay Figure S7

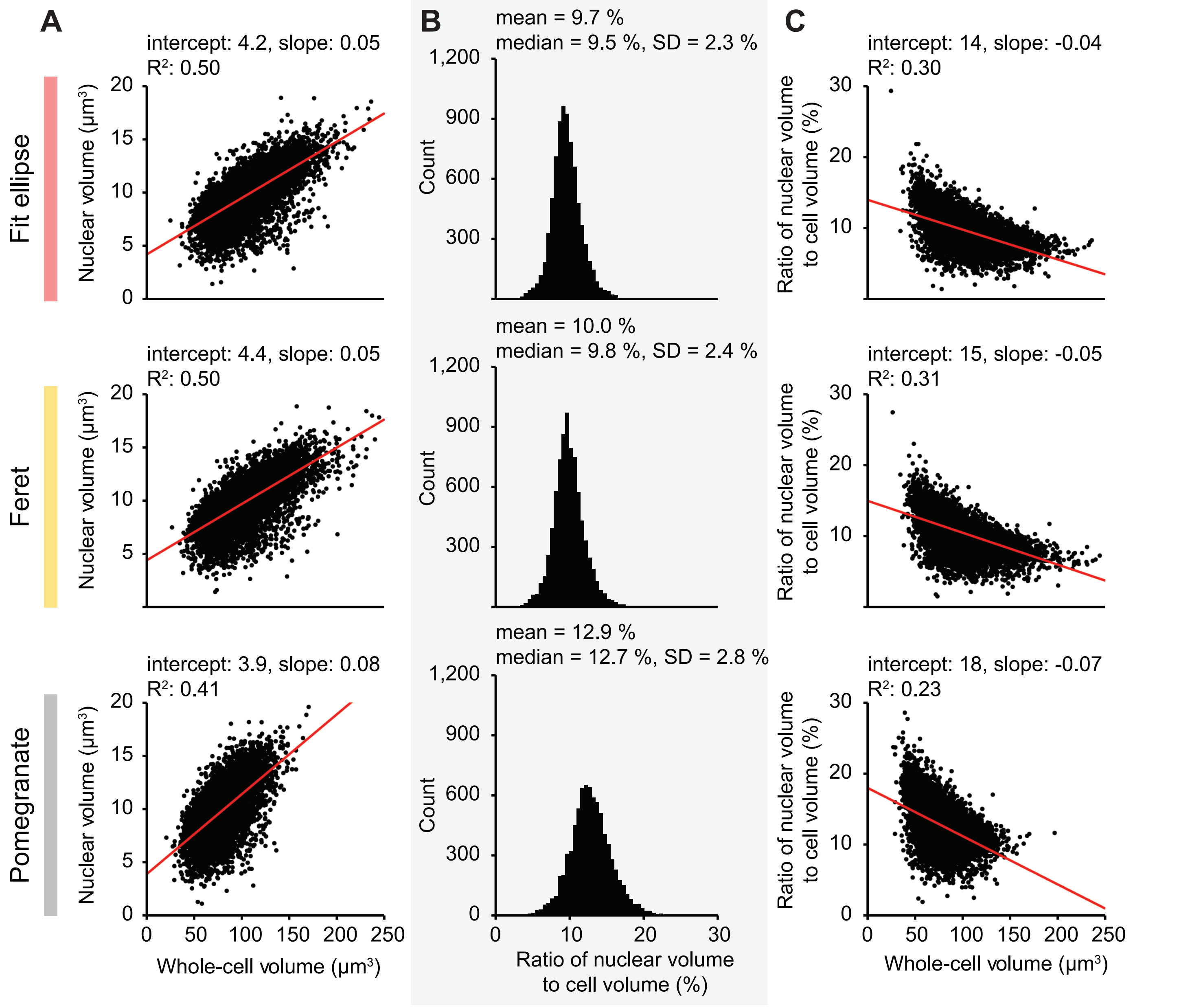


**Supplemental Figure S7. Wild-Type nuclear and cell volumes -** The same population of cells shown in S6, subject to both filters described in S2A (n = 8,026). **(A)** Nuclear and whole-cell volumes shown as scatterplots. Annotations provide information on the overlaid linear regression (red line). **(B)** Histograms of the ratio of nuclear volume to total cell volume (bins = 60). Ratios are calculated as the Pomegranate-reconstructed nuclear volumes divided by the whole-cell volumes derived from the various reconstruction methods. Annotations provide descriptive statistics (SD = standard deviation). **(C)** Scatterplots of the ratio of nuclear volume to total cell volume, compared to total cell volume. Annotations provide information on the overlaid linear regression (red line).

### Baybay Figure S8

**A**

| Group | Parameter | Cell density in image | Cell length (Feret diameter) | Cell length (fit ellipse) | Cell volume (Pomegranate) | Cell surface area (Pomegranate) | Cell width (Feret diameter) | Cell width (ellipsoidal fit) | Mean cell width (Pomegranate) |
|---|---|---|---|---|---|---|---|---|---|
| width | Mean cell width (Pomegranate) | −0.18 | × | × | 0.41 | 0.27 | 0.72 | 0.93 | |
| width | Cell width (ellipsoidal fit) | −0.17 | 0.17 | 0.15 | 0.56 | 0.43 | 0.89 | | 0.93 |
| width | Cell width (Feret diameter) | −0.11 | 0.28 | 0.25 | 0.60 | 0.49 | | 0.89 | 0.72 |
| size | Cell surface area (Pomegranate) | −0.11 | 0.96 | 0.95 | 0.99 | | 0.49 | 0.43 | 0.27 |
| size | Cell volume (Pomegranate) | −0.13 | 0.90 | 0.89 | | 0.99 | 0.60 | 0.56 | 0.41 |
| length | Cell length (fit ellipse) | −0.07 | 1 | | 0.89 | 0.95 | 0.25 | 0.15 | × |
| length | Cell length (Feret diameter) | −0.07 | | 1 | 0.90 | 0.96 | 0.28 | 0.17 | × |
| | Cell density in image | | −0.07 | −0.07 | −0.13 | −0.11 | −0.11 | −0.17 | −0.18 |

Correlation: 1.0, 0.5, 0.0, −0.5, −1.0

**B**

| | Cell density in image | Cell length (Feret diameter) | Cell length (fit ellipse) | Cell volume (Pomegranate) | Cell surface area (Pomegranate) | Cell width (Feret diameter) | Cell width (ellipsoidal fit) | Mean cell width (Pomegranate) |
|---|---|---|---|---|---|---|---|---|
| Mean cell width (Pomegranate) | < 0.0001 | 0.2107 | 0.6523 | < 0.0001 | < 0.0001 | < 0.0001 | < 0.0001 | |
| Cell width (ellipsoidal fit) | < 0.0001 | < 0.0001 | < 0.0001 | < 0.0001 | < 0.0001 | < 0.0001 | | < 0.0001 |
| Cell width (Feret diameter) | < 0.0001 | < 0.0001 | < 0.0001 | < 0.0001 | < 0.0001 | | < 0.0001 | < 0.0001 |
| Cell surface area (Pomegranate) | < 0.0001 | < 0.0001 | < 0.0001 | < 0.0001 | | < 0.0001 | < 0.0001 | < 0.0001 |
| Cell volume (Pomegranate) | < 0.0001 | < 0.0001 | < 0.0001 | | < 0.0001 | < 0.0001 | < 0.0001 | < 0.0001 |
| Cell length (fit ellipse) | < 0.0001 | < 0.0001 | | < 0.0001 | < 0.0001 | < 0.0001 | < 0.0001 | 0.6523 |
| Cell length (Feret diameter) | < 0.0001 | | < 0.0001 | < 0.0001 | < 0.0001 | < 0.0001 | < 0.0001 | 0.2107 |
| Cell density in image | | < 0.0001 | < 0.0001 | < 0.0001 | < 0.0001 | < 0.0001 | < 0.0001 | < 0.0001 |

**Supplemental Figure S8. Correlation of size parameters** - **(A)** Correlogram showing Pearson's r correlation coefficient on various combinations of size parameters, in addition to cell density in the image. Size parameters were extracted from the dataset presented in Figure S6 (n = 8,026). X denotes relationships with non-significant correlation ($p > 0.05$). Cell length correlates strongly with Pomegranate reconstruction volume, as well as with theoretical volumes - supporting the preexisting concept that cell length is a viable approximation for cell size. However, the additional positive correlation between cell width and cell volume illustrates the importance of taking cell width into account when approximating cell size. **(B)** Table of p-values for the correlations presented in (A). Red cells indicate non-significant correlations ($p > 0.05$).

### Baybay Figure S9

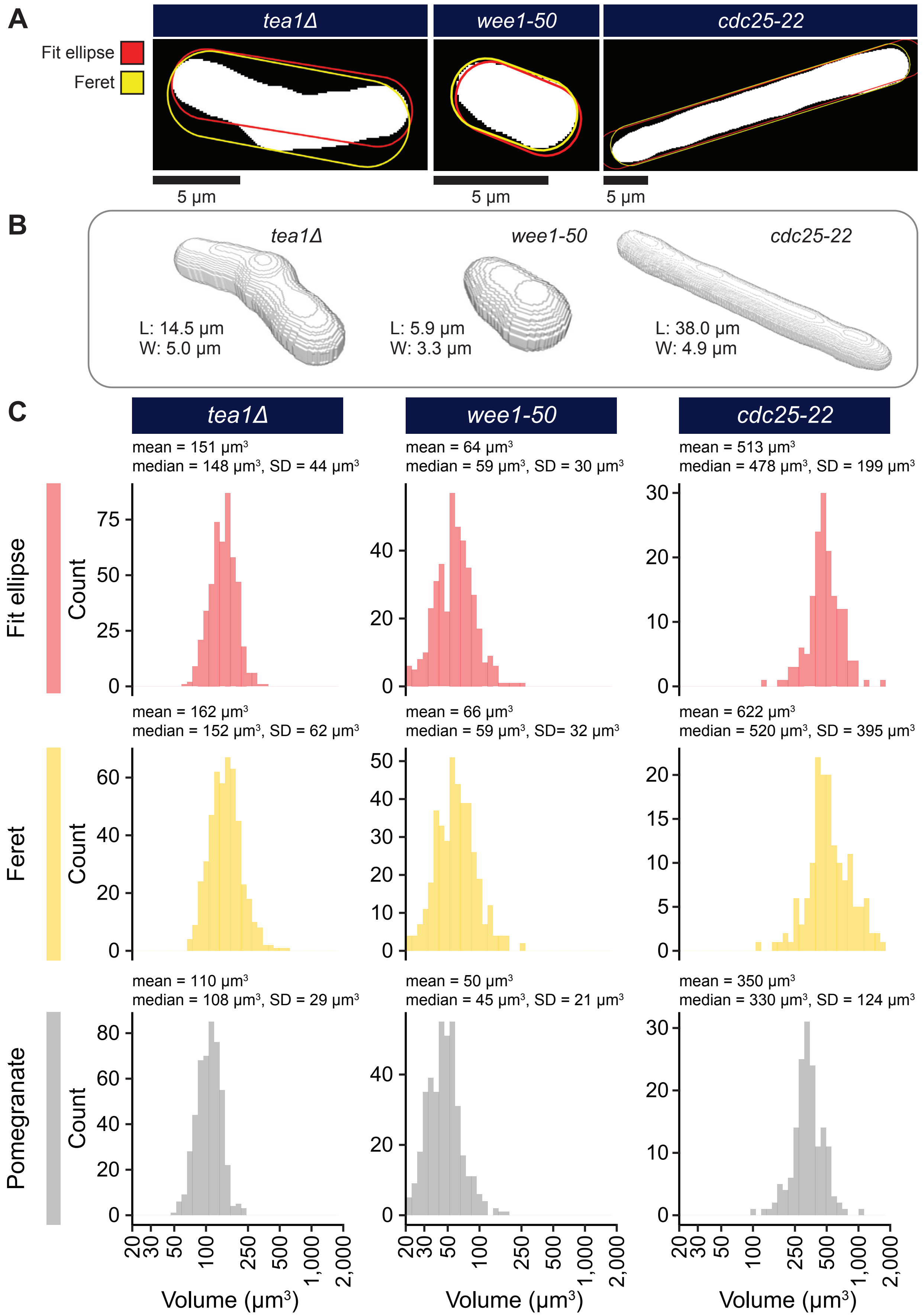


**Supplemental Figure S9. Comparison of methods for calculating cell volumes in size and shape mutants - (A)** Midplane ROI (white), with overlaid cross sections of the rod-shape obtained with length and width from ellipse fitting (red) or Feret diameters (yellow). **(B)** Reconstruction models of representative, individual cells. Voxel size is 0.1071 μm x 0.1071 μm x 0.065 μm. Annotations provide size parameters for each cell: maximum Feret diameter (length, L) and minimum Feret diameter (width, W). **(C)** Histograms of whole-cell volumes for size and shape mutants produced with various reconstruction methods as described in Figure 4. The x-axis of the volume histogram is on a base-10 logarithmic scale. Annotations provide descriptive statistics (SD = standard deviation, *tea1Δ*: n = 476 cells, *wee1-50*: n = 394 cells, *cdc25-22*: n = 157 cells).

### Baybay Figure S10

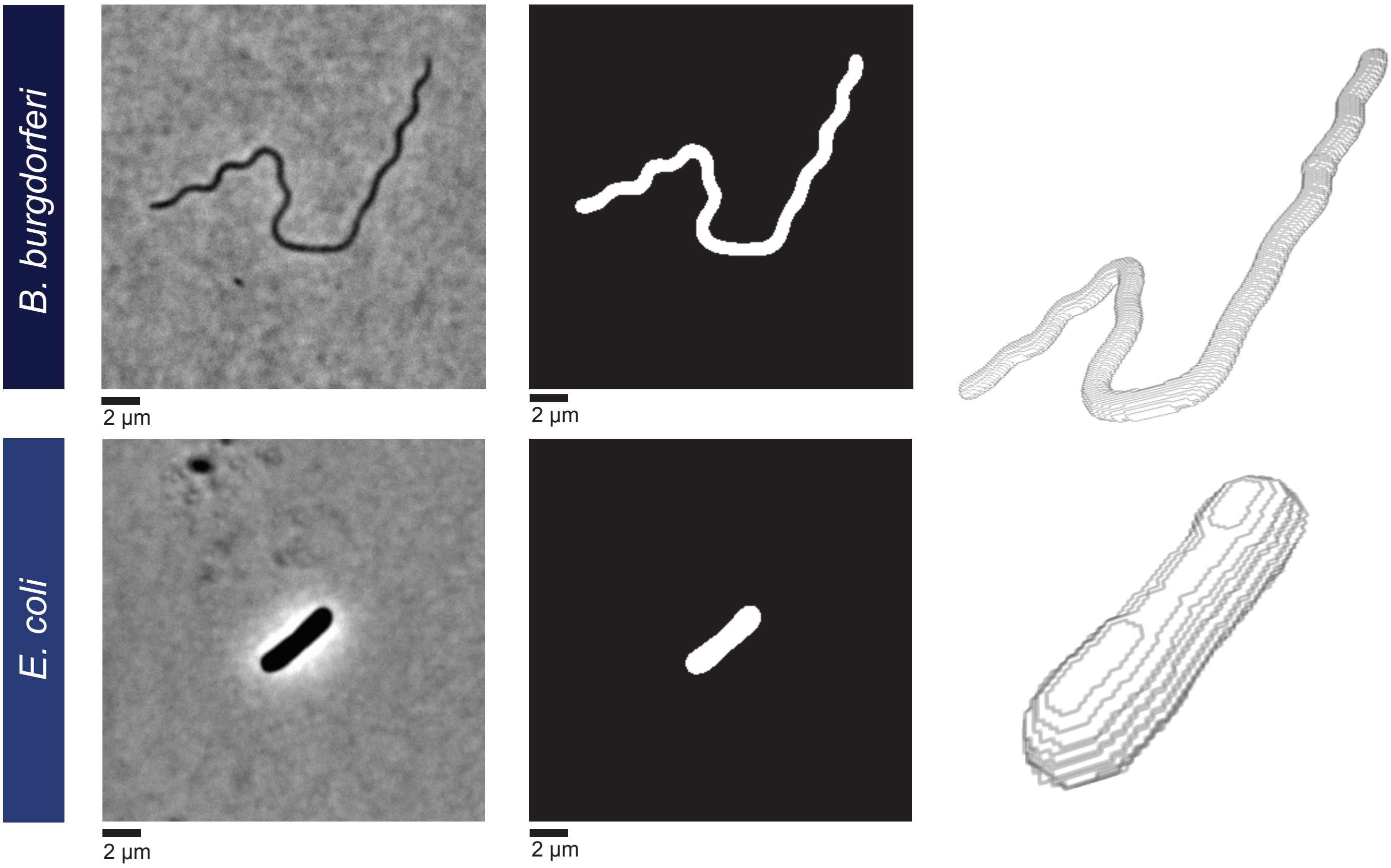


**Supplemental Figure S10. Sample reconstructions of bacteria -** Phase contrast images of *B. burgdorferi* and *E. coli*. (first column), binarized images using adaptive thresholding (second column), and Pomegranate reconstructions (third column). Voxel size is 0.064 µm x 0.064 µm x 0.064 µm. The phase contrast image is a single optical section; the voxel size in Z for the reconstruction is arbitrarily set to match that of the pixel size in XY. *E. coli* is rod-shaped (Koch, 2001; Cabeen and Jacobs-Wagner, 2005), whereas *B. burgdorferi* has been reported to have a planar waveform (Goldstein, Charon and Kreiling, 1994; Wolgemuth *et al.*, 2006). Pomegranate can only capture this wave accurately when it is oriented parallel to the imaging plane.


Cabeen, M. T. & Jacobs-Wagner, C. Bacterial cell shape. *Nat Rev Microbiol* 3, 601-610, doi:10.1038/nrmicro1205 (2005).

Goldstein, S. F., Charon, N. W. & Kreiling, J. A. Borrelia burgdorferi swims with a planar waveform similar to that of eukaryotic flagella. *Proc Natl Acad Sci U S A* 91, 3433-3437, doi:10.1073/pnas.91.8.3433 (1994).

Koch, A. L. in *Bacterial Growth and Form*, 271-330 (Springer Netherlands, 2001).

Wolgemuth, C. W., Charon, N. W., Goldstein, S. F. & Goldstein, R. E. The flagellar cytoskeleton of the spirochetes. *J Mol Microbiol Biotechnol* 11, 221-227, doi:10.1159/000094056 (2006).
